# Mathematical Details on a Cancer Resistance Model

**DOI:** 10.1101/475533

**Authors:** James M. Greene, Cynthia Sanchez-Tapia, Eduardo D. Sontag

## Abstract

The primary factor limiting the success of chemotherapy in cancer treatment is the phenomenon of drug resistance. We have recently introduced a framework for quantifying the effects of induced and non-induced resistance to cancer chemotherapy [11, 10]. In this work, we expound on the details relating to an optimal control problem outlined in [10]. The control structure is precisely characterized as a concatenation of bang-bang and path-constrained arcs via the Pontryagin Maximum Principle and differential Lie algebra techniques. A structural identifiability analysis is also presented, demonstrating that patient-specific parameters may be measured and thus utilized in the design of optimal therapies prior to the commencement of therapy. For completeness, a detailed analysis of existence results is also included.

## 1 Introduction

The ability of cancer chemotherapies to successfully eradicate cancer populations is limited by the presence of drug resistance. Cells may become resistant through a variety of cellular and micro-environmental mechanisms [9]. These mechanisms are exceedingly complex and diverse, and remain to be completely understood. Equally complex is the manner in which cancer cells obtain the resistant phenotype. Classically resistance was understood to be conferred by random genetic mutations; more recently, the role of epigenetic phenotype switching was discovered as a mediator of resistance as well [17]. Importantly, both of these phenomena are drug-independent, so that the generation of resistance is functionally separate from the selection mechanism (e.g. the drug). However, experimental studies from the past ten years suggest that drug resistance in cancer may actually be induced by the application of the chemotherapy [17, 21, 22, 8, 4].

In view of the multitude of ways by which a cancer cell may become chemoresistant, we have previously introduced a mathematical framework to differentiate both drug-independent and drug-dependent resistance [11]. In this work, we demonstrated that induced resistance may play a crucial role in therapy outcome, and also discussed methods by which a treatment’s mutagenic potential may be identified via biological assays. An extension of the work was presented in [10], where a formal optimal control problem was introduced, and an initial mathematical analysis was performed. The aim of this work is to both formalize the identifiability properties of our theoretical framework, establish the existence of the optimal control introduced in [10], and to precisely classify the optimal control structure utilizing the Pontryagin Maximum Principle and differential-geometric techniques. A numerical investigation of both the control structure and corresponding objective is also undertaken as a function of patient-specific parameters, and clinical conclusions are emphasized.

The work is organized as follows. In Section 2, we briefly review the mathematical model together with the underlying assumptions. Section 3 formalizes the structural identifability of model parameters. Section 4 restates the optimal control problem. Existence results are presented in Section 5. The Maximum Principle is analyzed in Section 6. A precise theoretical characterization of the optimal control structure is summarized in Section 7. In Section 8, we compare theoretical results with numerical computations, and investigate the variation in control structure and objective as a function of parameters. Lastly, conclusions are presented in Section 9.

## 2 Mathematical Modeling of Induced Drug Resistance

We briefly review the model presented in [11] and analyzed in [10]. In that work, we have constructed a simple dynamical model which describes the evolution of drug resistance through both drug-independent (e.g. random point mutations, gene amplification) and drug-dependent (e.g. mutagenicity, epigenetic modifications) mechanisms. To our knowledge, this is the first theoretical study of the phenomena of drug-induced resistance, which although experimentally observed remains poorly understood. It is our hope that a mathematical analysis will provide mechanistic insight and produce a more complete understanding of this process by which cancer cells inhibit treatment efficacy.

Specifically, we assume that the cancer population is composed of two types of cells: sensitive (*S*) and resistant (*R*). For simplicity, the drug is taken as completely ineffective against the resistant population, while the log-kill hypothesis [31] is assumed for the sensitive cells. Complete resistance is of course unrealistic, but can serve as a reasonable approximation, especially when toxicity constraints are considered, and hence limit the total amount of drug that may be administered. Furthermore, this assumption permits a natural metric on treatment efficacy that may not exist otherwise (see Section 4). The effect of treatment is considered as a control agent *u*(*t*), which we assume is a locally bounded Lebesgue measurable function taking values in ℝ_+_. Here *u*(*t*) is directly related to the applied drug dosage *D*(*t*), and in the present work we assume that we have explicit control over *u*(*t*). Later, during the formulation of the optimal control problem (Section 4), we will make precise specifications on the control set *U*. Obviously, a general measurable function may be unrealistic as a treatment strategy due to limitations in current clinical technologies. However our objective in this work is to understand the fundamental mathematical questions associated to drug-induced resistance. Furthermore, our results in Section 7 suggest that the applied optimal treatment should take a relatively simple form, which may be approximated with sufficient accuracy in a clinical setting.

Sensitive and resistant cells are assumed to compete for resources in the tumor microenvironment; this is modeled via a joint carrying capacity, which we have scaled to one. Furthermore, cells are allowed to transition between the two phenotypes in both a drug-independent and drug-dependent manner. All random transitions to the resistant phenotype are modeled utilizing a common term, *ϵS*, which accounts for both genetic mutations and epigenetic events occurring independently of the application of treatment. Drug-induced transitions are assumed of the form *αu*(*t*)*S*, which implies that the per-capita drug-induced transition rate is directly proportional to the dosage (as we assume full control on *u*(*t*), i.e. pharmacokinetics are ignored). Of course, other functional relationships may exist, but since the problem is not well-studied, we consider it reasonable to begin our analysis in this simple framework. The above assumptions then yield the following system of ordinary differential equations (ODEs):

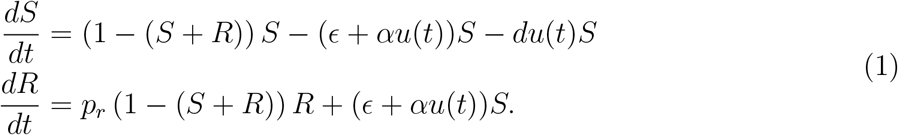

All parameters are taken as non-negative, and 0 ≤ *p_r_* < 1. The restriction on pr emerges due to (1) already being non-dimensionalized, as *p_r_* represents the relative growth rate of the resistant population with respect to that of the sensitive cells. The condition *p_r_* < 1 thus assumes that the resistant cells divide more slowly than their sensitive counterparts, which is both observed experimentally [14, 20, 3], and necessary for our mathematical framework. Indeed, the condition *p_r_* ≥ 1 would imply that *u*(*t*) ≡ 0 is optimal under any clinically realistic objective.

As mentioned previously, many simplifying assumptions are made in system (1). Specifically, both types of resistance (random genetic and epigenetic) are modeled as dynamically equivalent; both possess the same division rate *p_r_* and spontaneous (i.e. drug-independent) transition rate *ϵ*. Thus, the resistant compartment *R* denotes the total resistant subpopulation, both genetic and epigenetic.

The region Ω = {(*S, R*) | 0 ≤ *S* + *R* ≤ 1} in the first quadrant is forward invariant for any locally bounded Lebesgue measurable treatment function *u*(*t*) taking values in ℝ_+_. Furthermore, if *ϵ* > 0, the population of (1) becomes asymptotically resistant:

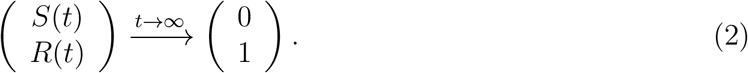

For a proof, see Theorem 2 in SI A in [11]. Thus, in our model, the phenomenon of drug resistance is inevitable. However, we may still implement control strategies which, for example, may increase patient survival time. Such aspects will inform the objective introduced in Section 4. For more details on the formulation and dynamics of system (1), we refer the reader to [11].

## 3 Structural Identifiability

Before beginning a discussion of the optimal control problem, we discuss the identifiability of system (1). Our focus in the remainder of the work is on control structures based on the presence of drug-induced resistance, and thus relies on the ability to determine whether, and to what degree, the specific chemotherapeutic treatment is generating resistance. Ideally, we envision a clinical scenario in which cancer cells from a patient are cultured in an *ex vivo* assay (for example, see [23]) prior to treatment. Parameter values are then calculated from treatment response dynamics in the assay, and an optimal therapy regime is implemented based on the theoretical work described below. Thus, identifying patient-specific model parameters, specially the induced-resistance rate *α*, is a necessary step in determining the control structures to apply. In this section, we address this issue, and prove that all parameters are structurally identifiable, as well as demonstrate a specific set of controls that may be utilized to determine *α*. A selfcontained discussion is presented; for more details on theoretical aspects, see [25] and the references therein. Other recent works related to identifiability in the biological sciences (as well as *practical identifiability*) can be found in [5, 32].

We first formulate our dynamical system, and specify the input and output variables. To mimic a cancer patient (and not, for instance, an in vivo experiment), the treatment *u*(*t*) is the sole input. Furthermore, we assume that the only clinically observable output is the entire tumor volume *V*(*t*):

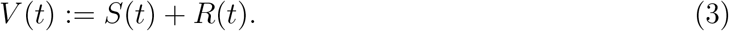

That is, we do not assume real-time measurements of the individual sensitive and resistant sub-populations. We note that in some instances, such measurements may be possible; however for a general chemotherapy, the precise resistance mechanism may be unknown *a priori*, and hence no biomarker with the ability to differentiate cell types may be available.

Treatment is initiated at time *t* = 0, at which we assume an entirely sensitive population:

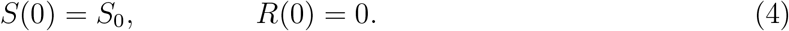

Here 0 < *S*_0_ < 1, so that (*S*(*t*), *R*(*t*)) ∈ Ω for all *t* ≥ 0. We note that *R*(0) = 0 is not restrictive, and similar results may derived under the more general assumption 0 ≤ *R*_0_ < 1. The condition *R*(0) = 0 is utilized both for computational simplicity and since *R*(0) is generally small (assuming a non-zero detection time, and small random mutation parameter *ϵ* see [11] for a discussion).

The above then allows us to formulate our system (1) in input/output form, where the input *u*(*t*) appears affinely:

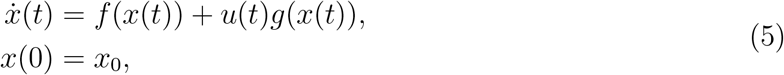

where *f* and *g* are

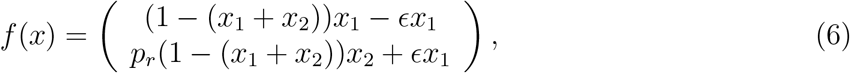

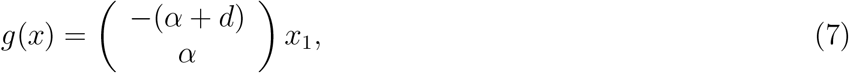

and *x*(*t*) = (*S*(*t*), *R*(*t*)). As is standard in control theory, the output is denoted by the variable *y*, which in this work corresponds to the total tumor volume:

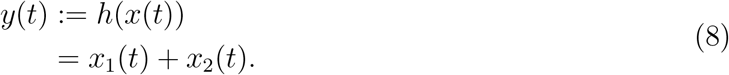

A system in form (5) is said to be *uniquely structurally identifiable* if the map *p* ↦ (*u*(*t*), *x*(*t, p*)) is injective almost everywhere [5, 16], where *p* is the vector of parameters to be identified. In this work,

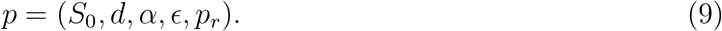

*Local identifiability* and *non-identifiability* correspond to the map being finite-to-one and infinite-to-one, respectively. Our objective is then to demonstrate unique structural identifiability for model system (5) (or equivalently (1)), and hence recover all parameter values *p* from only measurements of the tumor volume *y*. We also note that the notion of identifiability is closely related to that of *observability*; for details [1, 24] are a good reference.

To analyze identifiability, we utilize results appearing in [12, 33, 26], and hence frame the issue from a differential-geometric perspective. Our hypothesis is that perfect (hence noise-free) input-output data is available in the form of *y* and its derivatives on any interval of time. We thus, for example, make measurements of

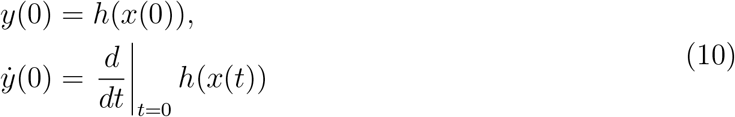

and relate their values to the unknown parameter values *p*. If there exist inputs *u*(*t*) such that the above system of equations may be solved for *p*, the system is identifiable. The right-hand sides of (10) may be computed in terms of the Lie derivatives of the vector fields *f* and *g* in system (5). We recall that Lie differentiation *L_X_ H* of a *C^ω^* function H by a *C^ω^* vector field *X*:

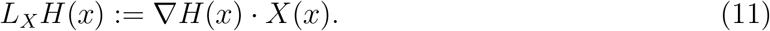

Here the domain of both *X* and *H* is the first-quadrant triangular region Ω, seen as a subset of the plane, and the vector fields and output function are *C^ω^* on an open set containing Ω (infact, they are given by polynomials, so they extend as analytic functions to the entire plane). Recall that set *C^ω^* consists of all analytic functions. Iterated Lie derivatives are well-defined, and should be interpreted as function composition, so that for example *L_Y_ L_X_H* = *L_Y_*(*L_X_H*), and 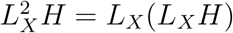.

More formally, defining observable quantities as the zero-time derivatives of the output *y* = *h*(*x*),

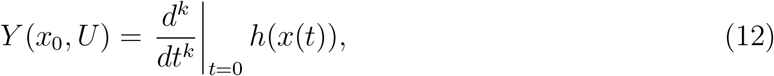

where *U* ∈ *R^k^* is the value of the control *u*(*t*) and its derivatives evaluated at *t* = 0: *U* = (*u*(0), *u′*(0),…, *u*^(*k*−1)^(0)). Here *k* ≥ 0, indicating that the *k*^th^-order derivative *Y* may expressed as a polynomial in the components of *U* [26]. The initial conditions *x*_0_ appear due to evaluation at *t* = 0. The observation space is then defined as the span of the *Y*(*x*_0_, *U*) elements:

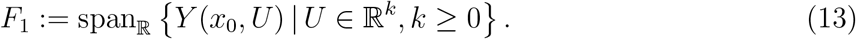

Conversely, we also define span of iterated Lie derivatives with respect to the output *h* and vector fields *f*(*x*) and *g*(*x*):

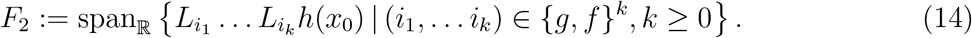

Wang and Sontag [33] proved that *F*_1_ = *F*_2_, so that the set of “elementary observables” may be considered as the set of all iterated Lie derivatives *F*_2_. Hence, identifiability may be formulated in terms of the reconstruction of parameters *p* from elements in *F*_2_. Parameters *p* are then identifiable if the map

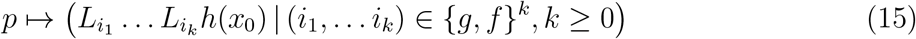

is one-to-one. For the remainder of this section, we investigate the mapping defined in (15).

Computing the Lie derivatives and recalling that *x*_0_ = (*S*_0_, 0) we can recursively determine the parameters *p*:

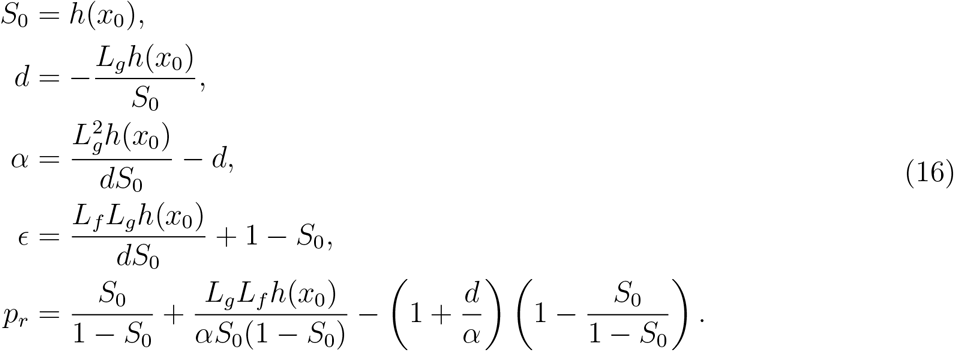

Since *F*_1_ = *F*_2_, all of the above Lie derivatives are observable via appropriate treatment protocols. For an explicit set of controls and corresponding relations to measurable quantities (elements of the form (12)), see [11]. Thus, we conclude that all parameters in system (1) are identifiable, which allows us to investigate optimal therapies dependent upon *a priori* knowledge of the drug-induced resistance rate *α*.

## 4 Optimal Control Formulation

As discussed in Section 2, all treatment strategies *u*(*t*) result in an entirely resistant tumor: (*S*_*_, *R*_*_) = (0, 1) is globally asymptotically stable for all initial conditions in region Ω. Thus, any chemotherapeutic protocol will eventually fail, and a new drug must be introduced (not modeled in this work, but the subject of future study). Therefore, selecting an objective which minimizes tumor volume (*S*+*R*) or resistant fraction (*R*/(*S*+*R*)) at a fixed time horizon would be specious for our modeling framework. However, one can still combine therapeutic efficacy and clonal competition to influence transient dynamics and possibly prolong patient life, as has been shown clinically utilizing real-time patient data [7]. Motivated by this observation, we define an objective based on maximizing time until treatment failure, as described below.

Let

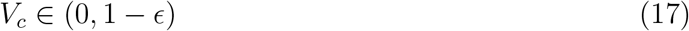

be a *critical tumor volume* at which treatment, by definition, has failed. The upper bound is a technical constraint that will be needed in Section 7; note that this is not prohibitive, as the genetic mutation rate *ϵ* is generally small [15], and our interest is on the impact of induced resistance. Recall that all cellular populations have been normalized to remain in [0, 1]. Our interpretation is that a tumor volume larger than *V_c_* interferes with normal biological function, while *S* + *R* ≤ *V_c_* indicates a clinically acceptable state. Different diseases will have different *V_c_* values. Define *t_c_* as the time at which the tumor increases above size *V_c_* for the first time. To be precise, *t_c_* is the maximal time for which *S* + *R* ≤ *V_c_*. Since all treatments approach the state (0, 1), *t_c_* is well defined for each treatment *u*(*t*): *t_c_* = *t_c_*(*u*). Time *t_c_* is then a measure of treatment efficacy, and our goal is then to determine *u*_*_ which maximizes *t_c_*(*u*).

Toxicity as well as pharmacokinetic constraints limit the amount of drug to be applied at any given instant. Thus, we assume that there exists *M* > 0 such that *u*(*t*) ≤ *M* for all *t* ≥ 0. Any other Lebesgue measurable treatment regime *u*(*t*) is then considered, so that the control set *U* = [0, *M*] and the set of admissible controls 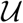 is

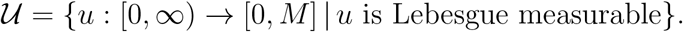

We are thus seeking a control 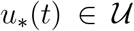 which maximizes *t_c_*, i.e. solves the time-optimal minimization problem

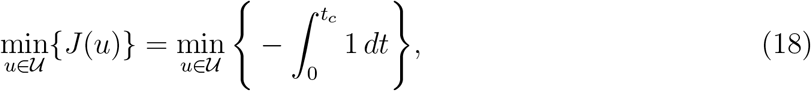

restricted to the dynamic state equations given by the system described previously in (5). Note that the above is formulated as a *minimization* problem to be consistent with previous literature and results related to the Pontryagin Maximum Principle (PMP) [13]. Note that maximization is still utilized in Sections 5 and 6.1, and we believe that the objective will be clear from context.

The time *t_c_* must satisfy the terminal condition (*t_c_, x*(*t_c_*)) ∈ *N*, where *N* is the line *S*+*R* = *V_c_* in Ω, i.e.

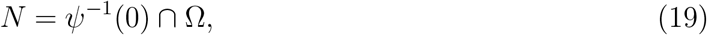

where

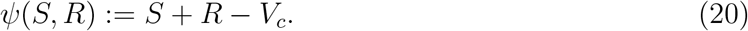

Furthermore, the path-constraint

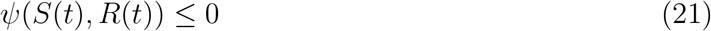

must also hold for 0 ≤ *t* ≤ *t_c_*. Equation (21) ensures that the tumor remains below critical volume *V_c_* for the duration of treatment. Equivalently, the dynamics are restricted to lie in the set Ω_*c*_ ⊆ Ω, where

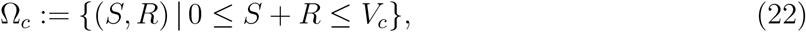

for times *t* such that *t* ∈ [0, *t_c_*]. The initial state

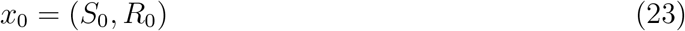

is also assumed to lie in Ω_*c*_. For the remainder of the work, we no longer restrict to the case *R*_0_ = 0 as was assumed for simplicity in Section 3.

## 5 Existence Results

Before characterizing the structure of the desired optimal control for the problem presented in Section 4, we must first verify that the supremum of times *t_c_*(*u*) for 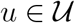 is obtained by some 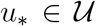, i.e. that the optimal control exists. This involves two distinct steps: proving that the supremum is both finite and that it is obtained by at least one admissible control. The following two subsections verify these claims.

### 5.1 Finiteness of the Supremum

We prove that

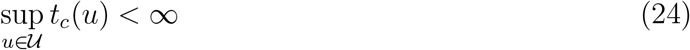

for a more general control system. The result depends crucially on (2), and the fact that asymptotically stable state (0, 1) is disjoint from the dynamic constraint *x* ∈ Ω_*c*_ (see equation (21)). That is, *V_c_* < 1 is necessary for the following subsequent result to hold, and generally an optimal control will not exist if *V_c_* = 1 or if the path constraint (21) is removed.

Consider a control system of the form

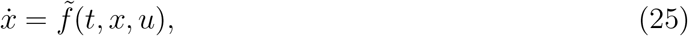

where *x* ∈ Ω, 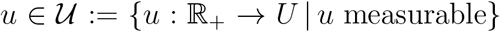, and 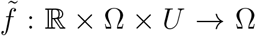, where Ω is an open subset of ℝ^*n*^, *U* ⊆ ℝ. Fix the initial conditions

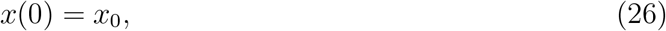

with *x*_0_ ∈ Ω, and assume that all solutions of (25) and (26) approach a fixed point 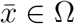. That is, for all 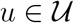,

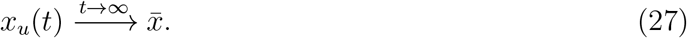

Note that we explicitly denote the dependence of the trajectory on the control *u*, and the above point 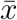 is independent of the control *u*.

Fix a closed subset *L* of Ω such that *x*_0_ ∈ *L*, 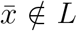. Associate to each control (and hence corresponding trajectory) a time *t_c_*(*u*) such that

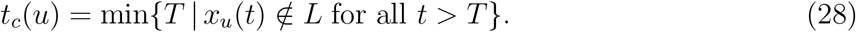

Note that the above is well-defined (as a minimum) for each control *u*, since by assumption *x*_0_ ∈ *L* and each trajectory asymptotically approaches 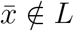.

#### Theorem 1.

*Define*

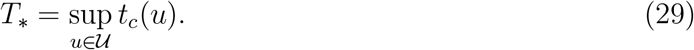

*With the above construction, T*_*_ *is finite*.

*Proof*. Fix an open set *K* of Ω containing 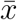 such that 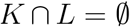. Note that this is possible, as *L* is closed and 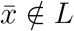. Suppose, by contradiction, that *T*_*_ = ∞. We construct a trajectory that remains in *L* for all time *t*, thus contradicting the fact that every trajectory must eventually enter *K*. By definition of the supremum, there exists a sequence of controls 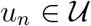 such that *x_u_n__* ∈ *L* for *t* ∈ [0, *t_n_*], with 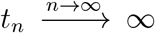. By taking a subsequence, we assume that *t_n_* is increasing. Note that each *u_n_* is defined on [0, ∞), and the initial condition (26) is fixed for all pairs (*x_u_n__, u_n_*).

We construct a new control *u*_*_ inductively as follows. Define a sequence of controls on [0, *t*_1_]:

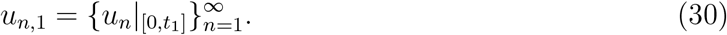

Since the time interval [0, *t*_1_] is compact, there exists a convergence subsequence 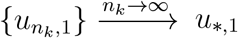 in the weak topology. As a subsequence, the corresponding trajectory *x*_*,1_ remains in *L* for *t* ∈ [0, *t*_1_]. Similarly, define subsequence *u*_*n*,2_ on [*t*_1_, *t*_2_] as

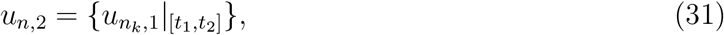

where we begin the sequence at the maximum of *n*_1_ and 2, to ensure that all controls in the sequence correspond to trajectories entirely inside of *L*. Again, since the interval is compact, there exists a convergence subsequence (again in the weak topology) of *u*_*n*,2_, say *u*_*,2_, with, by continuity of the state with respect to the control, the corresponding trajectory lies entirely in *L*. Continue in this manner, constructing a sequence of controls *u*_*,*i*_, *i* = 1, 2,…, where *u*_*,*i*_ is defined on [*t*_*i*−1_, *t_i_*], with *t*_0_ = 0. Define *u*_*_ on [0, ∞) as the concatenation of the *u*_*,*i*_:

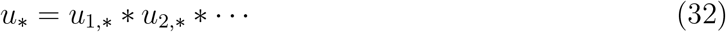

As the pointwise limit of measurable functions, *u*_*_ is measurable. Clearly, *u* ∈ *U* by construction, so that 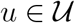. The corresponding trajectory *x*_*u*_*__ thus lies entirely in *L* for *t* ∈ [0, ∞), and hence never enters *K*. This the desired contradiction, so that *T*_*_ must be finite, as desired.

#### Proof option 2 for Theorem 1

Consider the sets *K, V* ⊂ ℝ^2^, with *V* being an open neighborhood of the steady state (0, 1) and *K* a compact set in ℝ^2^ such that

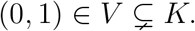

Suppose, by contradiction, that *T*_*_ is not finite, so we can find a sequence of controls 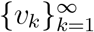 in 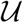 satisfying

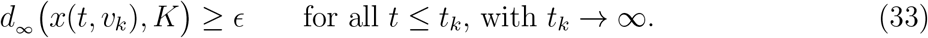

where *d*_∞_ denotes the supremum metric and, for each 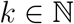, *x*(*t, v_k_*) is the solution of the IVP:

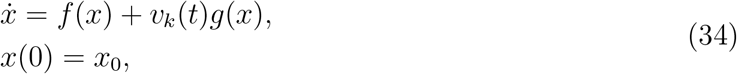

Our aim is to find a control 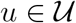 such that *x*(*t, u*), solution of system (34), does not enter *K* for any *t* > 0. In order to do so, we make us of the Banach-Alaoglu’s theorem that guarantees that the ball

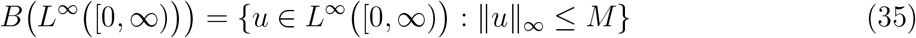

is a compact set on the weak^*^ topology and metrizable. Thus, the sequence 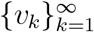 must have a weak^*^–convergent subsequence 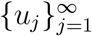 which converges to a control *u* ∈ *L*^∞^ ([0, ∞)). In other words, for every *ψ* ∈ *L*^1^ ([0, ∞))

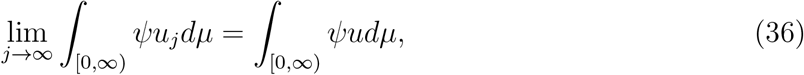

where *μ* is the usual Lebesgue measure. This means that the sequence 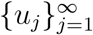 converges to *u* with respect to the weak^*^ topology on *L*^∞^ [0, ∞) as the dual of *L*^1^ [0, ∞).

We next prove that lim_*j*→∞_ ║*x*(*t,u*) – *x*(*t, u_j_*)║_∞_ = 0 for all *t* ∈ [*t*_*k*−1_, *t_k_*] and all *k* ∈ ℕ. In order to do so define

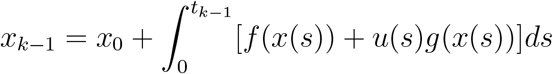

for any *t*_*k*−1_ ∈ [0, ∞), where *x* solves the IVP

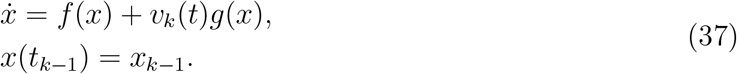

Notice that the fact that the equilibrium (0, 1) is globally asymptotically stable implies that *x*_*k*−1_ is well defined for any *k* ∈ ℕ.

The integral form of (37) is given by

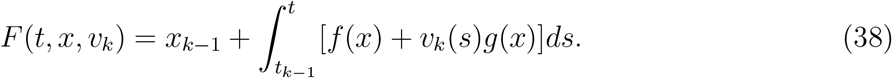

With the help of the *t_k_*’s from (33) and assuming *t_k_* ↑ ∞ as *k* goes to infinity we write the set [0, ∞) as the countable union of finite closed intervals:

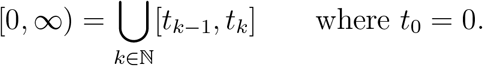

For each *k* ∈ ℕ, denote by 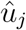 and 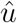 to the restricted functions of *u_j_* and *u* to the interval [*t*_*k*™1_, *t_k_*].

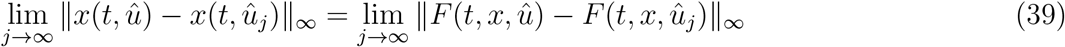

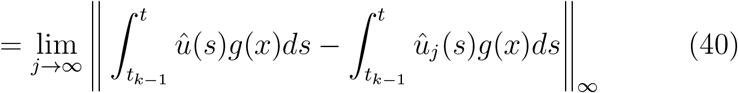

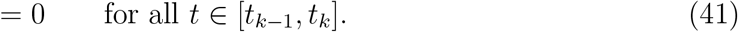

Since this result is independent of *k*, this implies that, for all *t* ∈ [*t*_*k*−1_, *t_k_*] and all *k* ∈ ℕ

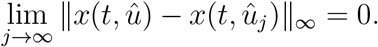

Thus,

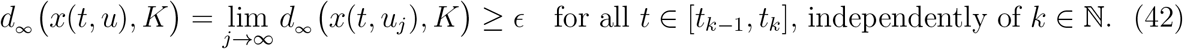

Therefore, the supremum of *t_c_* must be finite.

For the system and control problem defined in Sections 2 and 4, the above theorem implies that 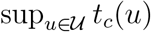 is finite by taking *L* = Ω_*c*_.

### 5.2 Supremum as a Maximum

Here we provide a general proof for the existence of optimal controls for systems of the form (5), assuming the set of maximal times is bounded above, which we have proven for our system in Section 5.1. For convenience, we make the proof as self-contained as possible (one well-known result of Filippov will be cited), and state the results in generality which we later apply to the model of induced resistance. Arguments are adapted primarily from the textbook of Bressan and Piccoli [2].

Consider again general control systems as in Section 5.1. Solutions (or trajectories) of (25) will be defined as absolutely continuous functions for which a control 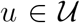 exists such that (*x*(*t*), *u*(*t*)) satisfy (25) a.e. in their (common) domain [*a, b*].

It is easier and classical to formulate existence with respect to differential inclusions. That is, define the multi-function

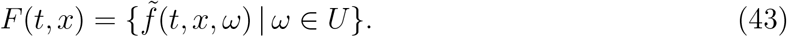

Thus, the control system (25) is clearly related to the inclusion

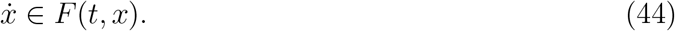

The following theorem (see [6] for a proof) makes this relationship precise.

#### Theorem 2.

*An absolutely continuous function x*: [*a, b*] → ℝ^*n*^ *is a solution of* (25) *if and only if it satisfies* (44) *almost everywhere*.

We first prove a lemma demonstrating that the set of trajectories is closed with respect to the sup-norm ║·║_∞_ if the set of velocities *F*(*t,x*) are all convex.

#### Lemma 3.

*Let x_k_* be a sequence of solutions of (25) *converging to x uniformly on* [0, *T*]. *If the graph of* (*t*, *x*(*t*)) *is entirely contained in* Ω, *and the F*(*t, x*) *are all convex, then x is also a solution of* (25).

*Proof*. By the assumptions on 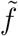, the sets *F*(*t,x*) are uniformly bounded as (*t,x*) range in a compact domain, so that *x_k_* are uniformly Lipschitz, and hence *x* is Lipschitz as the uniform limit. Thus *x* is differentiable a.e., and by Theorem 2, it is enough to show that

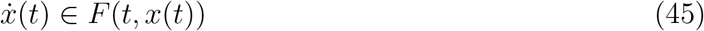

for all *t* such that the derivative exists.

Assume not, i.e. that the derivative exists at some *τ*, but 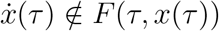. Since *F*(*τ, x*(*τ*)) is compact and convex, and 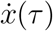 is closed, the Hyperplane Separation Theorem implies that there exists a hyperplane separating *F*(*τ, x*(*τ*)) and 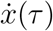. That is, there exists an *ϵ* > 0 and a (WLOG) unit-vector *p* ∈ ℝ^*n*^ such that

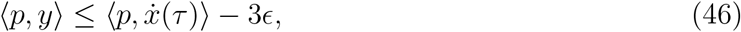

for all *y* ∈ *F*(*τ, x*(*τ*)). By continuity, there exists *δ* > 0 such that for |*t* – *t′*| ≤ *δ*, |*x′* – *x*(*τ*)| ≤ *δ*

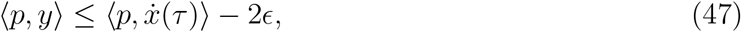

for all *y* ∈ *F*(*τ, x′*). Since *x* is differentiable at *τ*, we can choose *τ′* ∈ (*τ, τ* + *δ*] such that

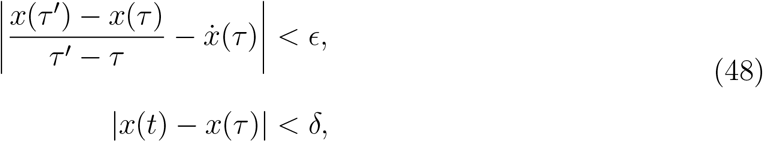

for all *t* ∈ [*τ, τ′*]. Equation (48) and uniform convergence then implies that

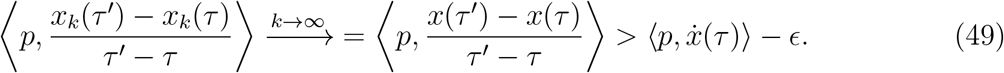

On the other hand, since 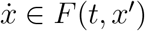 for *t* ∈ [*τ, τ′*], equation (47) implies that for *k* sufficiently large,

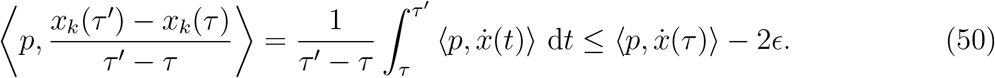

Clearly, (49) and (50) contradict one another, so that (45) must be true, as desired.

An optimal control problem associated to (25) may now be formulated. Suppose *x*_0_ ∈ Ω_*c*_ is an initial condition for the corresponding system, so that

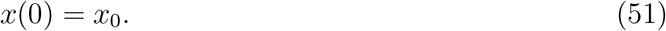

Let *S* denote the set of admissible terminal conditions, *S* ⊂ ℝ × ℝ^*n*^, and *ϕ*: ℝ × ℝ^*n*^ → ℝ a cost function. We would like to maximize *ϕ*(*T, x*(*T*)) over admissible controls with initial and terminal constraints:

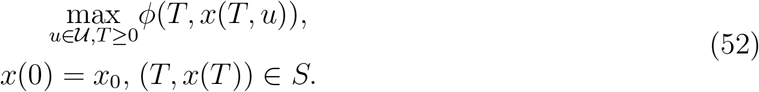

We now state sufficient conditions for such an optimal control to exist.

#### Theorem 4.

*Consider the control system* (25) *and corresponding optimal control problem* (52). *Assume the following*:

1. *The objective ϕ is continuous*.
2. *The sets of velocities F*(*t, x*) *are convex*.
3. *The trajectories x remain uniformly bounded*.
4. *The target set S is closed*.
5. *A trajectory satisfying the constraints in* (52) *exists*.
6. *S is contained in some strip* [0, *T*] × ℝ^*n*^, *i.e. the set of final times* (*for free-endpoint problems*) *can be uniformly bounded*.

*If the above items are al l satisfied, an optimal control exists*.

*Proof*. By assumption, there is at least one admissible trajectory reaching the target set *S*. Thus, we can construct a sequence of controls *u_k_*: [0, *T_k_*] → *U* whose corresponding trajectories *x_k_* satisfy

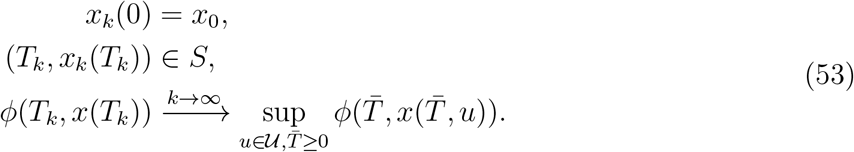

Since *S* ⊂ [0, *T*] × ℝ^*n*^, we know that *T_k_* ≤ *T* for all *k*. Each function *x_k_* can then be extended to the entire interval [0, *T*] by setting *x_k_*(*t*) = *x_k_*(*T_k_*) for *t* ∈ [*T_k_, T*].

The sequence *x_k_* is uniformly Lipschitz continuous, as *f* is uniformly bounded on bounded sets. This then implies equicontinuity of 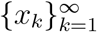. By the Arzela-Ascoli Theorem, there exists a subsequence *x_n_k__* such that *T_n_k__* → *T*_*_, *T*_*_ ≤ *T*, and *x_n_k__* → *x*_*_ uniformly on [0, *T*_*_].

Lemma 3 implies that *x*_*_ is admissible, so that there exists a control *u*_*_: [0, *T*_*_] ↦ *U* such that

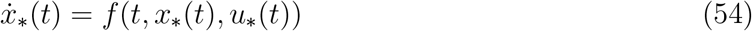

for almost all *t* ∈ [0, *T*_*_]. Equations (53) imply that

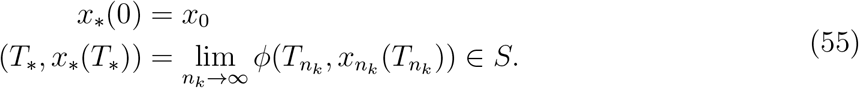

Note that the second of (55) relies on *S* being closed. Continuity of *ϕ* and (53) implies that

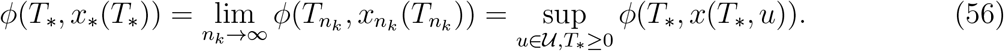

Thus, *u*_*_ is optimal, as desired.

For the model of drug-induced resistance, the right-hand side 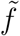 takes the form (5) where *f* and *g* are smooth on the domain Ω. Here the control set *U* is the compact set *U* = [0, *M*], and for such control-affine systems, convexity of *F*(*t, x*) is implied by the convexity of *U*. Existence of a trajectory satisfying the constraints is clear; for example, take *U*(*t*) ≡ 0. Our objective is to maximize the time to not escape the set *N*. Note that *N* is a closed subset of ℝ^2^, and that

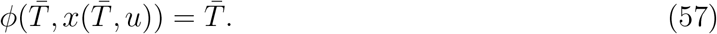

is continuous. Lastly, we have seen that all solutions remain in the closure 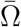, so that |*x*(*t*)| ≤ 1 for all 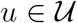 and hence solutions are uniformly bounded. Existence is then reduced to Item 6 in the previous theorem. Since the supremum was shown to be finite, Theorem 4 together with Theorem 1 imply that the optimal control for the problem presented in Section 4 exists.

## 6 Maximum Principle

The results of Section 5 imply that the optimal control problem introduced in Section 4 has at least one solution 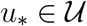. We now characterize this control utilizing the Pontryagin Maximum Principle (PMP). We envision a clinical scenario in which cancer cells from a patient are cultured in an *ex vivo* assay (for example, see [23]) prior to treatment. Parameter values are then calculated from treatment response dynamics in the assay, and an optimal therapy regime is implemented based on the theoretical work of this section. Thus, identifying patient-specific model parameters, specially the induced-resistance rate *α*, is a necessary step in determining control structures to apply. This issue was addressed partially in Section 3; for further *in vitro* results, see [11]. Hence, for the remainder of this work, we assume that prior to the onset of treatment, all patient-specific parameters are known. We now analyze behavior and response of system (1) to applied treatment strategies *u*(*t*) utilizing geometric methods. The subsequent analysis is strongly influenced by the Lie-derivative techniques introduced by Sussmann [27, 29, 28, 30]. For an excellent source on both the general theory and applications to cancer biology, see the textbooks by Schättler and Ledzewicz [13, 19].

### 6.1 Elimination of Path Constraints

We begin our analysis by separating interior controls from those determined by the path-constraint (21) (equivalently, *x* ∈ *N*). The following theorem implies that outside of the manifold *N*, the optimal pair (*x*_*_, *u*_*_) solves the same local optimization problem without the path and terminal constraints. More precisely, the necessary conditions of the PMP (see Section 6.2) at states not on N are exactly the conditions of the corresponding maximization problem with no path or terminal constraints.

#### Theorem 5.

*Suppose that x_*_ is an optimal trajectory. Let t*_1_ *be the first time such that x*(*t*) ∈ *N*. Fix ϵ > 0 *such that t*_1_ – *ϵ* > 0, *and*

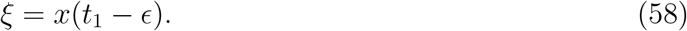

*Define z*(*t*):= *x*_*_(*t*)|_*t*∈[0,*t*_1_−*ϵ*]_. *Then the trajectory z is a local solution of the corresponding time maximization problem t_c_ with boundary conditions x*(0) = *x*_0_, *x*(*t_c_*) = *ξ, and no additional path constraints. Hence at all times t, z (together with the corresponding control and adjoint) must satisfy the corresponding unconstrained Pontryagin Maximum Principle*.

*Proof*. We first claim that *z* satisfies the path-constrained maximization problem with boundary conditions *x*(0) = *x*_0_, *x*(*t_c_*) = *ξ*. Otherwise, if there exists a trajectory 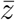 such that 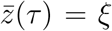, *τ* > *t*_1_ – *ϵ*, concatenate 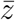 with *x*_*_ at *t* = *τ* to obtain a feasible trajectory satisfying all constraints. This trajectory then has total time *τ* + *ε* + *t_c_* – *t*_1_ > *t_c_*, contradicting the global optimality of *x*_*_.

Recall that *t*_1_ was the first time that *x*_*_(*t*) ∈ *N*. Since *z* is compact, we can find a neighborhood of *z* that lies entirely in {*x*| *x* ∉ *N*}. As the Maximum Principle is a local condition with respect to the state, this completes the proof.

Theorem 5 then tells us that for states *x* = (*S, R*) such that *S* + *R* < *V_c_*, the corresponding unconstrained PMP must be satisfied by any extremal lift of the original problem. Furthemore, the constraint (21) has generic order one. In other words,

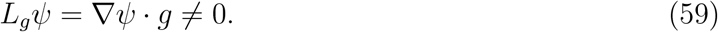

Therefore, the feedback control (also known as the constrained control) can be found by differentiating function (20) for trajectories to remain on the line *N*:

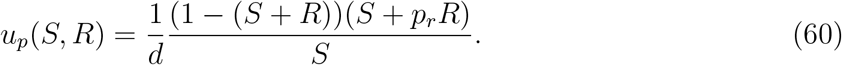

Its existence however, does not imply its feasibility, which is discussed below. Specifically, *u_p_* can be shown to a decreasing function of *S* which is feasible on the portion of *N* satisfying 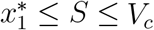, where 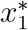 is given in (61), and infeasible elsewhere. This is proven in Proposition 7, and the geometric structure is depicted in Figure 1. Propositions 8 and 9 provide characterizations on the volume dynamics in certain regions of phase space, and are included here for completeness.

**Figure 1:**
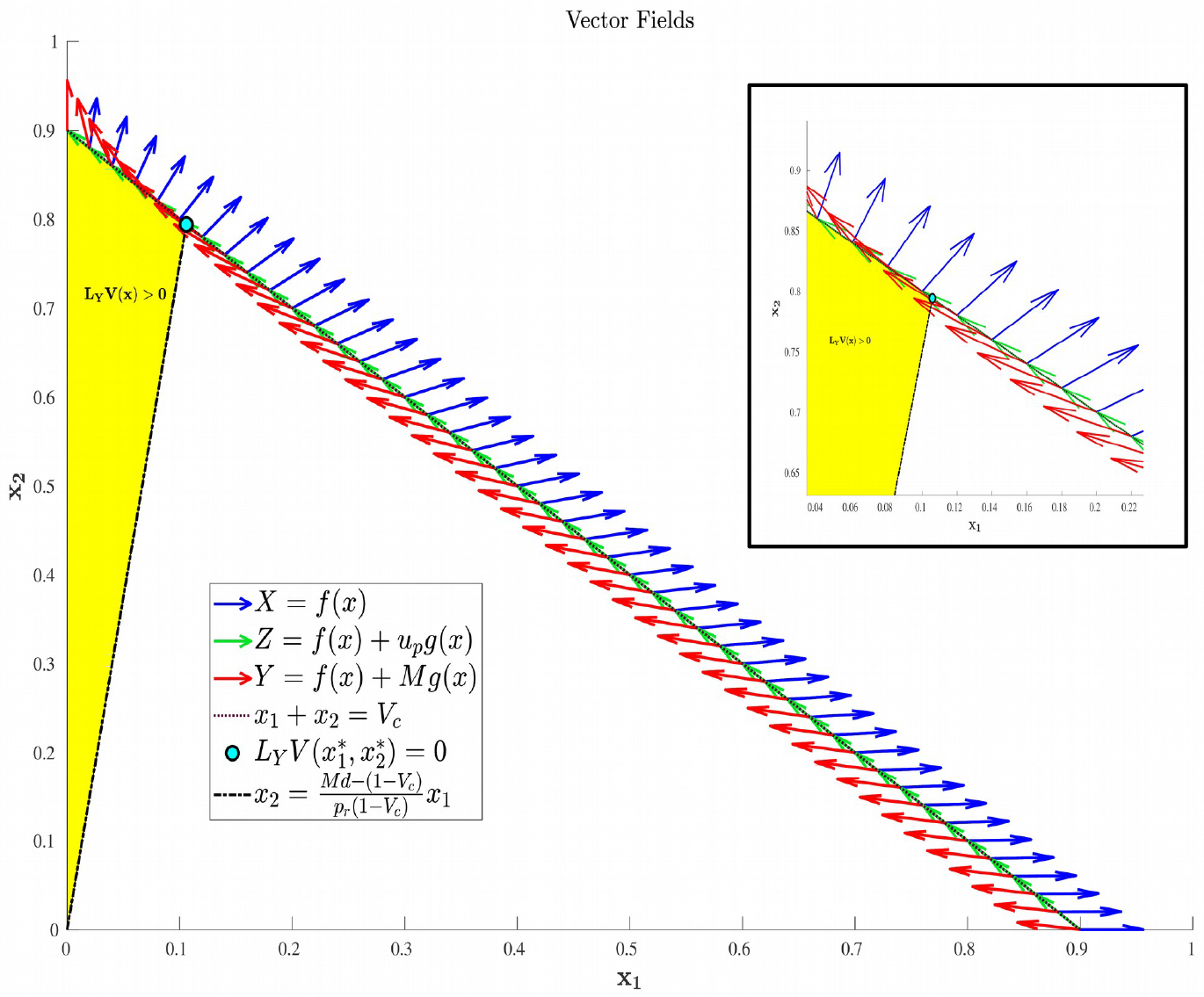
Region in Ω_*c*_ where *L_Y_V*(*x*) is guaranteed to be positive. That is, applying the maximal allowed dosage *M* causes the cancer population to increase in size for points in the yellow region. The figure is provided for visual clarity of the result in Proposition 8.

#### Proposition 6.

*Consider the volume function V*(*x*) = *S*+*R, and the point x*^*^ = (*S*^*^, *R*^*^) ∈ *N with coordinates*

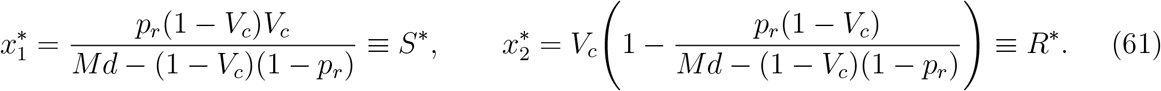

*Denote by y the vector field corresponding to the maximal allowed dosage M. The Lie derivative L_Y_V*(*x*) *of V*(*x*) *with respect to y is*

a. *positive if* 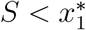,
b. *zero at* 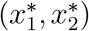, *and*
c. *negative if* 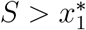.

*where* 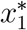 *is well defined (and positive) if*

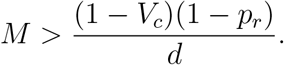

*Proof*. We verify the above claims with a direct calculation. For *x* ∈ *N*,

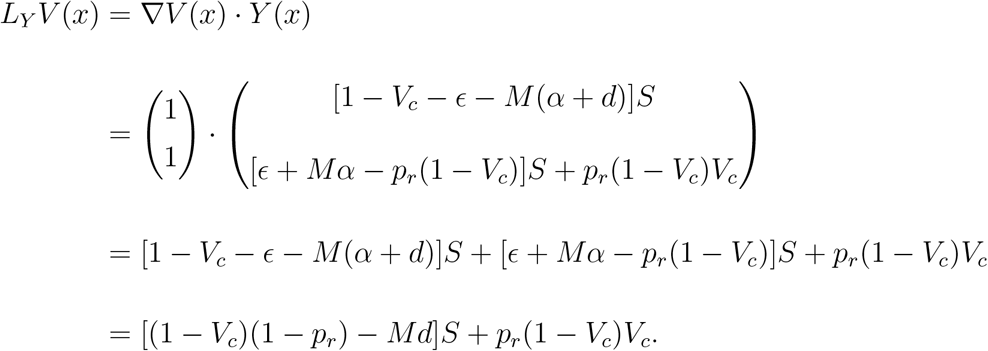

Assuming 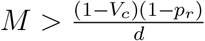, the sign of *L_Y_V*(*x*) is given as in the statement of the proposition.

#### Proposition 7.

*Let x be a point on the line N. The feedback control u_p_ is unfeasible if* 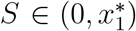 *and, it is feasible if* 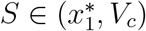

*Proof*. For *x* ∈ *N* we compute

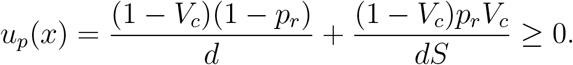

it is east to check that *u_p_* > *M* if 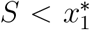. In addition, the feedback control, when restricted to points in *N*, is a decreasing function with respect to *S*. Thus, it is feasible for *x* ∈ *N* if 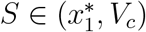.

#### Proposition 8.

*For x* = (*S, R*) ∈ Ω_*c*_ *with*

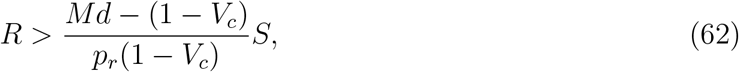

*the Lie derivative L_Y_V*(*x*) *is positive*.

*Proof*. Again, we compute *L_Y_V*(*x*):

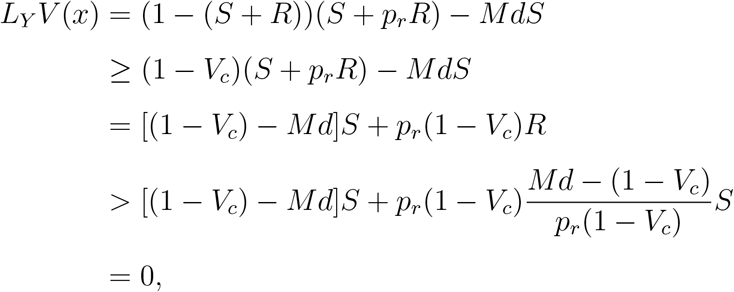

where the first inequality utilizes *V* ≤ *V_c_*, and the second relies on (62)

#### Proposition 9.

*For*

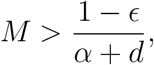

*trajectories corresponding to the maximal dosage M have a decreasing sensitive cellular population*.

*Proof*. for *u*(*t*) ≡ *M*, the corresponding sensitive trajectory is given by

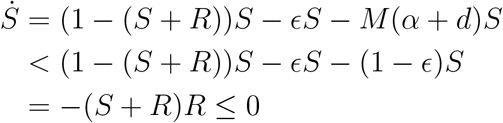

Note that we are assuming here that the maximal dosage *M* satisfies 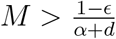.

We have now demonstrated that the optimal control consists of concatenations of controls obtained from the unconstrained necessary conditions and controls of the form (60). In the next section, we analyze the Maximum Principle in the region *S* + *R* < *V_c_*.

### 6.2 Maximum Principle and Necessary Conditions at Interior Points

Necessary conditions for the optimization problem discussed in Section 4 without path or terminal constraints are derived from the Pontryagin Maximum Principle [18, 13]. The corresponding Hamiltonian function *H* is defined as

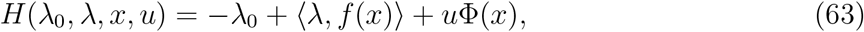

where *λ*_0_ ≥ 0 and *λ* ∈ ℝ^2^. Here 〉·, ·〈 denotes the standard inner product on ℝ^2^ and, since the dynamics are affine in the control *u*, Φ(*x*, λ) is the switching function:

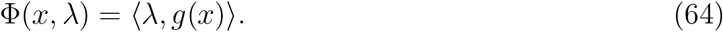

The Maximum Principle then yields the following theorem:

#### Theorem 10.

*If the extremal* (*x*_*_, *u*_*_) *is optimal, there exists* λ_0_ ≥ 0 *and a covector (adjoint)* λ: [0, *t_c_*] → (ℝ^2^)^*^, *such that the following hold:*

1. (λ_0_, λ(*t*)) ≠ 0 *for all t* ∈ [0, *t_c_*].
2. λ(*t*) = (λ_*S*_(*t*), λ_*R*_(*t*)) *satisfies the second-order differential equation*

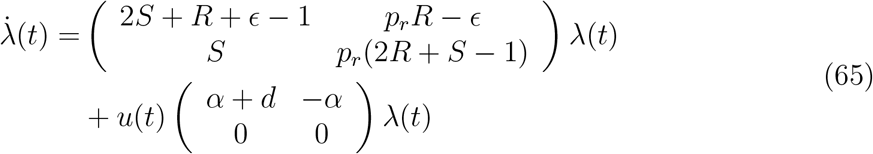
3. *u*_*_(*t*) *minimizes H pointwise over the control set U:*

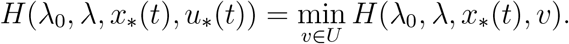 *Thus, the control u*_*_(*t*) *must satisfy*

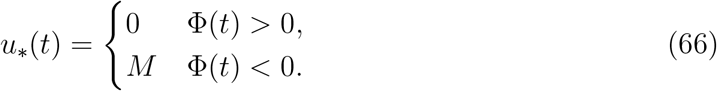

*where*

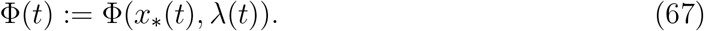
4. *The Hamiltonian H is identically zero along the extremal lift* (*x*_*_(*t*), *u*_*_(*t*), λ(*t*)):

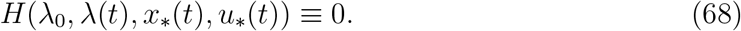

*Proof*. Most statements of Theorem 10 follow directly from the Maximum Principle, so proofs are omitted. In particular, items (1), (2) and the first part of (3) are immediate consequences [13]. Equation (66) follows directly since we minimize the function *H*, which is affine in *u* (see equation (63)). The Hamiltonian vanishes along (*x*_*_(*t*), *u*_*_(*t*), λ(*t*)) since it is independent of an explicit time *t* dependence and the final time *t_c_* is free, the latter being a consequence of the transversality condition.

For completeness, we state the following proposition.

#### Proposition 11.

*For all t* ∈ [0, *t_c_*], *the adjoint* λ(*t*) *corresponding to the extremal lift* (*x*_*_(*t*), *u*_*_(*t*), λ(*t*)) *is nonzero*.

*Proof*. This is a general result relating to free end time problems. We include a proof here for completeness. Suppose that there exists a time *t* ∈ [0, *t_c_*] such that λ(*t*) = 0. By (63), the corresponding value of the Hamiltonian is *H*(λ_0_, λ(*t*), *x*_*_(*t*), *u*^*^(*t*)) = –λ_0_. By item (4) in Theorem 10, *H* ≡ 0, which implies that λ_0_ = 0. This contradicts item (1) in Theorem 10. Hence, λ(*t*) ≠ 0 on [0, *t_c_*].

### 6.3 Geometric Properties and Existence of Singular Arcs

We now undertake a geometric analysis of the optimal control problem utilizing the affine structure of system (5) for interior states (i.e. controls which satisfy Theorem 10). We call such controls *interior extremals*, and all extremals in this section are assumed to be interior. The following results depend on the independence of the vector fields *f* and *g*, which we use to both classify the control structure for abnormal extremal lifts (extremal lifts with λ_0_ = 0), as well as characterize the switching function dynamics via the Lie bracket.

#### Proposition 12.

*For all S* ∈ Ω, *S* > 0, *the vector fields f*(*x*) *and g*(*x*) *are linearly independent*.

*Proof*. Define *A*(*x*) = *A*(*S, R*) to be the matrix

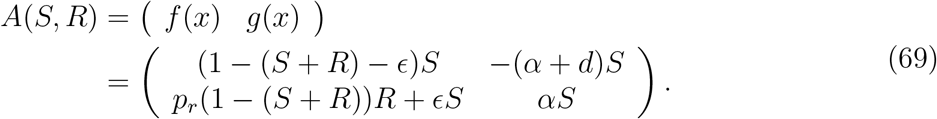

The determinant of *A* can calculated as

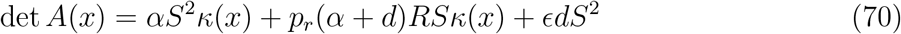

where

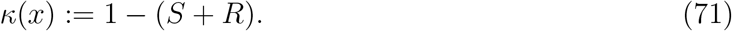

As *S*(*t*) + *R*(*t*) ≤ 1 for all *t* ≥ 0, *κ*(*x*(*t*)) ≥ 0, and we see that det *A*(*x*) = 0 in Ω if and only if *S* = 0, completing the proof.

The line *S* = 0 is invariant in Ω, and furthermore the dynamics in the set are independent of the control *u*(*t*). Conversely, *S*_0_ > 0 implies that *S*(*t*) > 0 for all *t* ≥ 0. We concern our analysis only in this latter case, and so without loss of generality, **f**(**x**) **and g**(**x**) **are linearly independent in the region of interest Ω_c_**.

We begin by showing that abnormal extremal lifts are easily characterized. We recall that an extremal lift is abnormal if λ_0_ = 0, i.e. if the Hamiltonian is independent of the objective.

#### Theorem 13.

*Abnormal extremal lifts at interior points, i.e. extremal lifts corresponding to* λ_0_ = 0, *are constant and given by the maximal* (*M*) *or minimal* (0) *dosage*.

*Proof*. Assume that *u*_*_ switches values at some time *t*. From (66), we must have that Φ(*t*) = 0. Since λ_0_ = 0 and Φ(*t*) = (λ(*t*), *g*(*x*_*_(*t*))), equation (63) reduces to

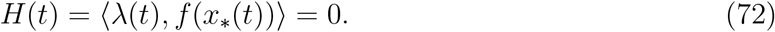

Thus, λ(*t*) is orthogonal to both *f*(*x*_*_(*t*)) and *g*(*x*_*_(*t*)). Since *f* and *g* are linearly independent (Proposition 12), this implies that λ(*t*) = 0. But this contradicts Proposition 11. Hence, no such time *t* exists, and *u*_*_(*t*) is constant. The constant sign of Φ thus corresponds to *u* = 0 or *u* = *M* (see equation (66)).

The control structure for abnormal extremal lifts is then completely understood via Theorem 13. To analyze the corresponding behavior for normal extremal lifts, without loss of generality we assume that λ_0_ = 1. Indeed, λ(*t*) may be rescaled by λ_0_ > 0 to yield an equivalent version of Theorem 10. We thus assume that the Hamiltonian *H*(*t*) evaluated along (λ(*t*), *x*_*_(*t*), *u*_*_(*t*)) is of the form

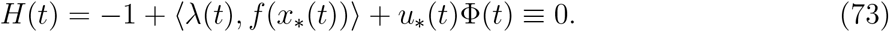

We recall the Lie bracket as the first-order differential operator between two vector fields *X*_1_ and *X*_2_:

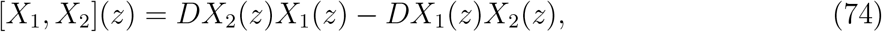

where, for example, *DX*_2_(*z*) denotes the Jacobian of *X*_2_ evaluated at *z*. As *f* and *g* are linearly independent in Ω, there exist *γ, β* ∈ *C*^∞^(Ω) such that

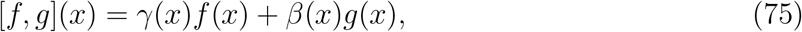

for all *x* ∈ Ω. In fact, we can compute *γ* and *β* explicitly:

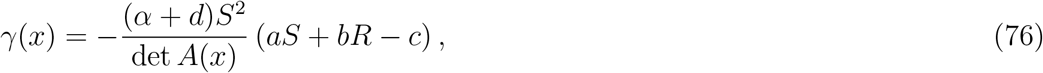

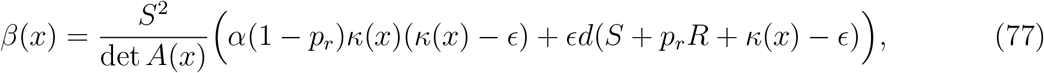

where

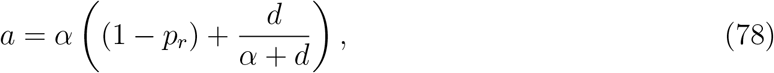

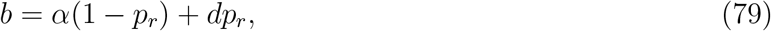

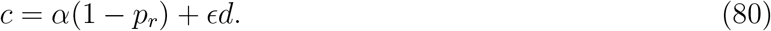

Clearly, for parameter values of interest (recall 0 < *p_r_* < 1), *a, b, c* > 0. The assumption (17) guarantees that *β*(*x*) > 0 on 0 < *S* + *R* < *V_c_*.

From (66), the sign of the switching function Φ determines the value of the control *u*_*_. As λ and *x*_*_ are solutions of differential equations, Φ is differentiable. The dynamics of Φ can be understood in terms of the Lie bracket [*f, g*]:

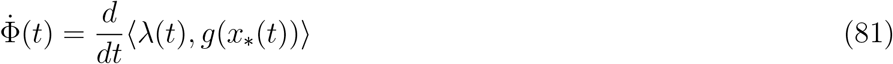

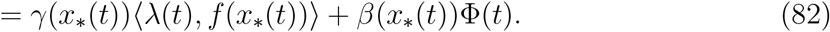

The last lines of the above follow from (75) as well as the linearity of the inner product. We are then able to derive an ODE system for *x*_*_ and Φ. Equation (73) allows us to solve for 〈λ(*t*), *f*(*x*_*_(*t*))〉:

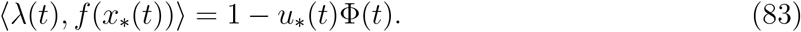

Substituting the above into (82) then yields the following ODE for Φ(*t*), which we view as coupled to system (5) via (66):

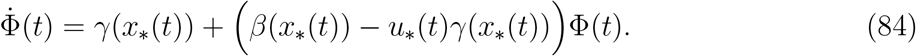

The structure of the optimal control at interior points may now be characterized as a combination of bang-bang and singular arcs. We recall that the control (or, more precisely, the extremal lift) *u*_*_ is singular on an open interval *I* ⊂ [0, *t_c_*] if the switching function Φ(*t*) and all its derivatives are identically zero on *I*. On such intervals, equation (66) does not determine the value of *u*_*_, and a more thorough analysis of the zero set of Φ(*t*) is necessary. Indeed, for a problem such as ours, aside from controls determined by the path constraint *ψ*(*S*(*t*), *R*(*t*)) ≤ 0, singular arcs are the only candidates for optimal controls that may take values outside of the set {0, *M*}. Conversely, times *t* where Φ(*t*) = 0 but Φ^(*n*)^(*t*) ≠ 0 for some *n* ≥ 1 denote candidate bang-bang junctions, where the control may switch between the vertices 0 and *M* of the control set *U*. Note that the parity of the smallest such n determines whether a switch actually occurs: n odd implies a switch, while for *n* even *u*_*_ remains constant. Equation (84) allows us to completely characterize the regions in the (*S, R*) plane where singular arcs are attainable, as demonstrated in the following proposition.

#### Proposition 14.

*Singular arcs are only possible in regions of the* (*S, R*) *plane where γ*(*x*) = 0. *Furthermore, as S*(*t*) > 0 *for all t* ≥ 0, *the region* {*x* ∈ ∝^2^ | *γ*(*x*) = 0} ⋂ Ω *is the line*

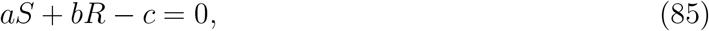

*where a, b, c are defined in* (78)–(80).

*Proof*. As discussed prior to the statement of Proposition 14, a singular arc must occur on a region where both Φ(*t*) and 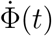 are identically zero (as well as all higher-order derivatives). Denoting by *x*_*_(*t*) the corresponding trajectory in the (*S, R*) phase plane, we may calculate 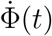 from equation (84):

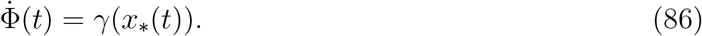

Note we have substituted the assumption Φ(*t*) = 0. Clearly we must also have that *γ*(*x*_*_(*t*)) = 0, thus implying that *x*_*_(*t*) ∈ *γ*^−1^(0), as desired. The last statement of the proposition follows immediately from equation (76).

Proposition 14 implies that singular solutions can only occur along the line *aS* + *bR* – *c* = 0. Thus, define regions in the first quadrant as follows:

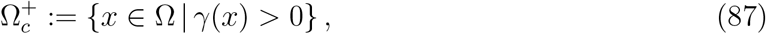

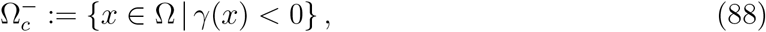

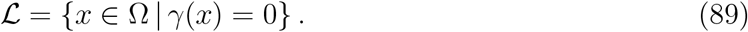

Recall that Ω_*c*_ is simply the region in Ω prior to treatment failure, i.e. 0 ≤ *V* ≤ *V_c_*. From (76), Ω_*c*_ is partitioned as in Figure 2(a). From (76) and (78)-(80), 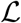 is a line with negative slope –*b*/*a*. Furthermore, necessary and sufficient conditions for 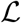 to lie interior to Ω_0_ are 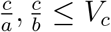. From (78)-(80), this occurs if and only if

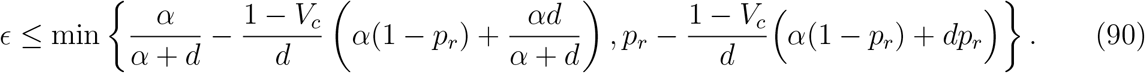

As *ϵ* is generally assumed small (recall that it represents the drug-independent mutation rate) and *V_c_* ≈ 1, this inequality is not restrictive, and we assume it is satisfied for the remainder of the work. We note an important exception below: when *α* = 0 the inequality is never satisfied with *ϵ* > 0; for such parameter values, line 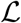 is horizontal. We note that this does not change the qualitative results presented below. Of course, other configurations of the line *aS* + *bR* = *c* and hence precise optimal syntheses may exist, but we believe the situation illustrated in Figure 2(a) is sufficiently generic for present purposes.

With the existence of singular arcs restricted to the line *γ* = 0 by Proposition 85, we now investigate the feasibility of such solutions. Recall that the treatment *u*(*t*) must lie in the control set *U* = [0, *M*], for some *M* > 0 corresponding to the maximally tolerated applied dosage. Defining the vector field *X*(*x*) and *Y*(*x*) as the vector fields corresponding to the vertices of *U*,

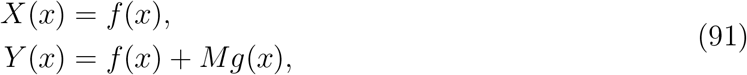

a singular control takes values in *U* at 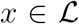 if and only if *X*(*x*) and *Y*(*y*) point in different directions along 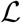. More precisely, the corresponding Lie derivatives *L_Xγ_*(*x*) and *L_Yγ_*(*x*) must have opposite signs; see Figure 2(b). The following proposition determines parameter values where this occurs.

**Figure 2:**
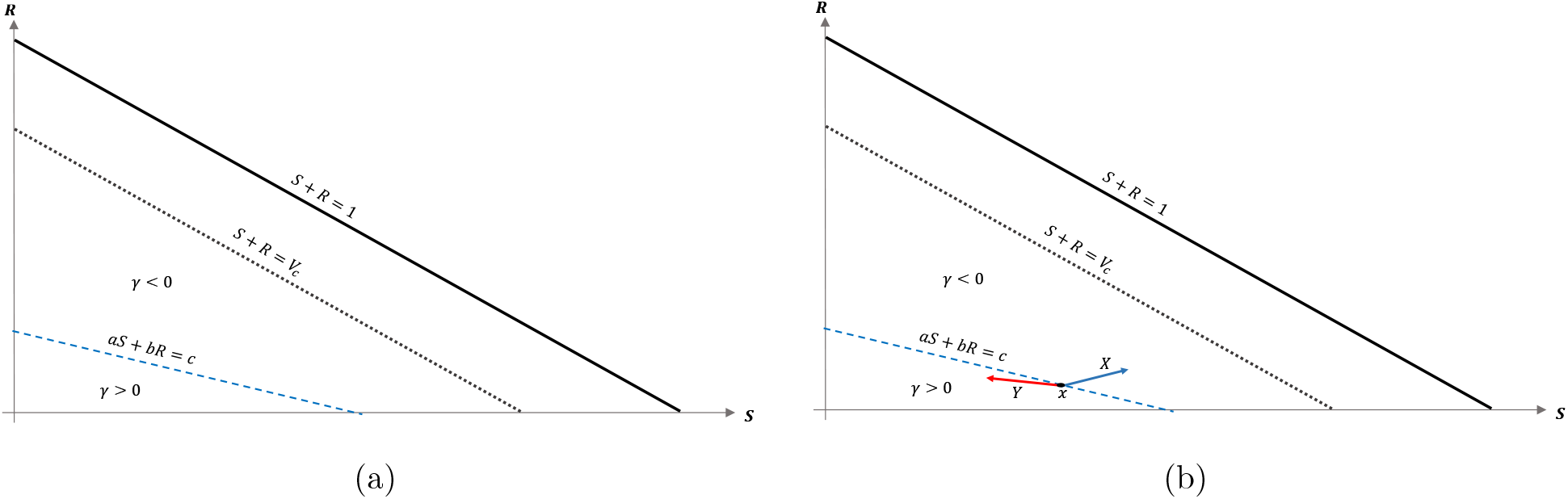
Domain in (*S, R*) plane. (a) Region where *γ* changes sign. We see that inside the triangular region *S* + *R* ≤ 1 of the first quadrant, *γ* changes sign only along the line *aS* + *bR* — *c* = 0. For this line to be interior to Ω_*c*_ as depicted, we must be in the parameter regime indicated in (90). (b) *X* and *Y* vector fields corresponding to vertices of control set *U*. For singular controls to lie in *U, X* and *Y* must point to opposite sides along 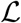.

#### Proposition 15.

*Suppose that α* > 0, *so that drug has the potential to induce drug resistance. Also, let the maximally tolerated dosage M satisfy*

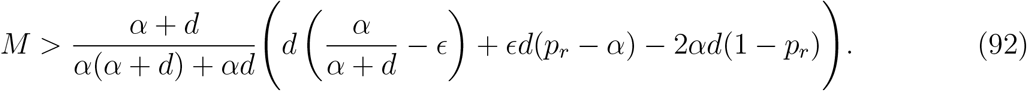

*Then the following hold along* 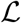:

1. *L_Xγ_* < 0,
2. *L_Yγ_* < 0 *as* 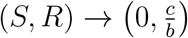 *in* Ω_0_,
3. *L_Yγ_* > 0 *at* 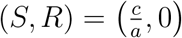, *and*
4. *L_Yγ_ is monotonically decreasing as a function of S*.

*Thus*, 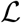 *contains a segment* 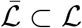 *which is a singular arc. Note that* 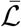 *is precisely the region in* 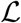 *where L_Yγ_ is negative*.

*Proof*. The proof is purely computational.

The geometry of Proposition 15 is illustrated in Figure 3. Thus, assuming *α* > 0 and *M* as in (92), singular arcs exist along the segment 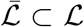. Furthermore, the corresponding control has a unique solution *u_s_*, which may be computed explicitly. Indeed, as the solution must remain on the line 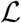, or equivalently, *aS* + *bR* = *c*, taking the time derivative of this equation yields 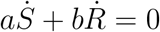, and substituting the expressions (1) we compute *u_s_* as

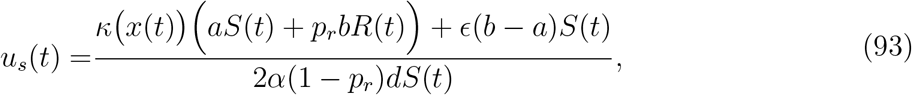

where *a, b, c* are given by (78)-(80) and *R* and *S* satisfy *aS* + *bR* = *c*. As discussed previously, *S*(*t*) > 0 for *S*_0_ > 0, so this formula is well-defined. Proposition 15 implies that it is possible to simplify equation (93) as a function of *S* (i.e. as a *feedback law*) for 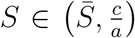, for some 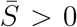, but since its value will not be needed, we do not provide its explicit form. Note that the maximal dose *M* is achieved precisely at 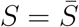 where vector field *Y* is parallel to 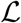. Thus, at this 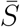, the trajectory must leave the singular arc, and enter the region 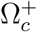. As 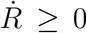, trajectories must follow 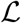 in the direction of decreasing *S*; see Figure 3. We summarize these results in the following theorem.

**Figure 3:**
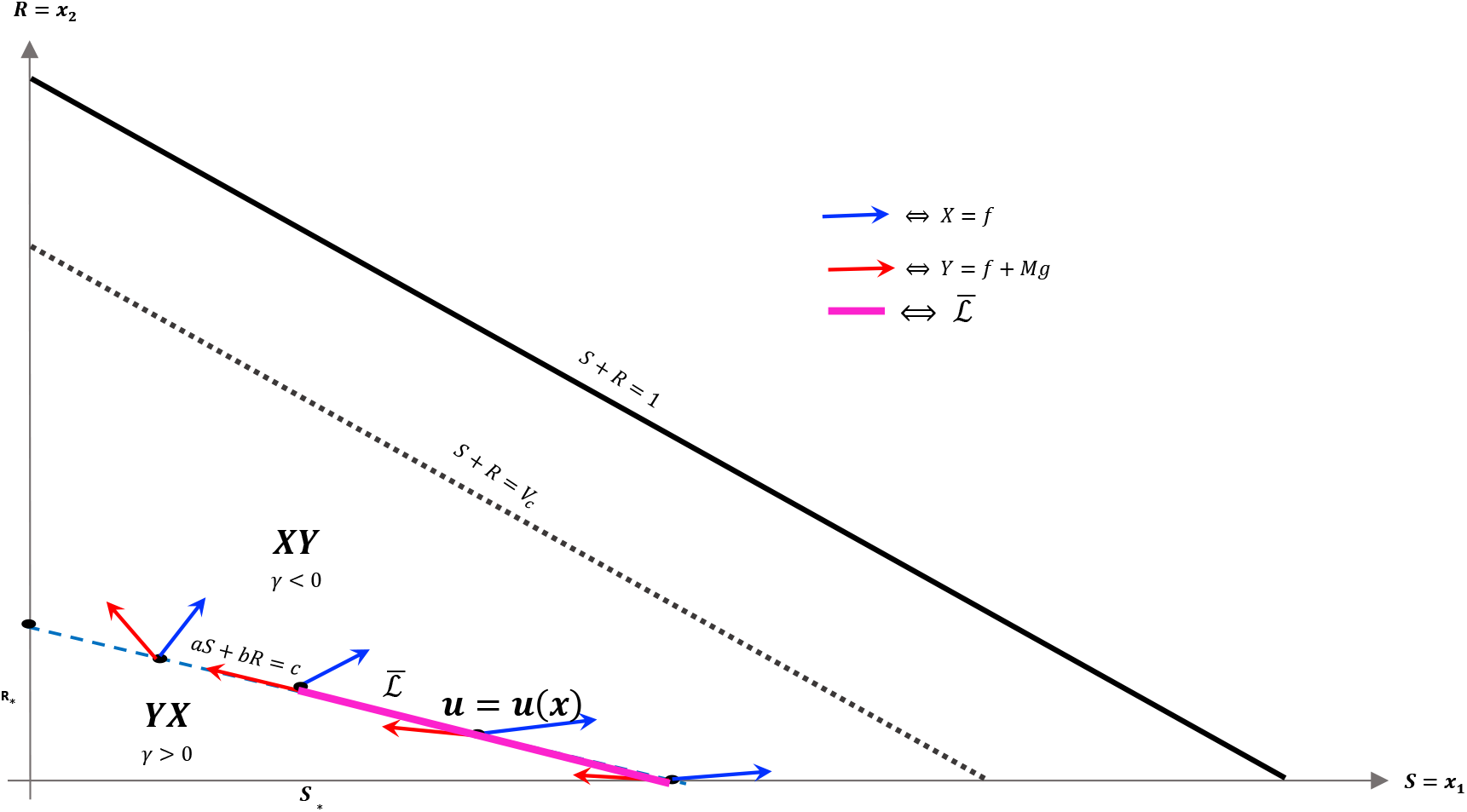
Geometry of vector fields *X* and *Y* with *α* > 0 and *M* satisfying (92). As in Proposition 15, this can be understood via the corresponding Lie derivatives of *γ*. Note that near *R* = 0, *X* and *Y* point to opposite sides of 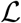, while at 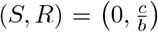, both *X* and *Y* point away from *γ* > 0. The line 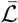 is the unique singular arc in Ω_*c*_.

#### Theorem 16.

*If α* > 0, *and M satisfies* (92), *a singular arc exists in the* (*S, R*) *plane as a segment of the line* 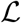. *Along this singular arc, the control is given by equation* (93), *where aS* + *bR* = *c. Therefore, in this case the necessary minimum conditions on u*_*_ *from* (66) *can be updated as follows:*

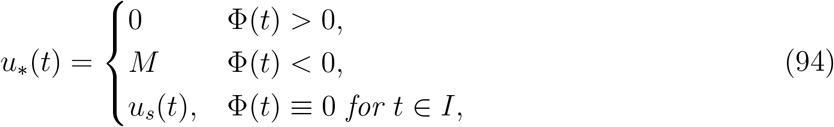

*where I is an open interval. Recall again that this is the optimal control at points interior to* Ω_*c*_.

*Proof*. See the discussion immediately preceding Theorem 16.

In the case *α* = 0, the line 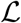 is horizontal, and as *R* is increasing, no segment 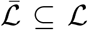 is admissible in phase space. Thus, the interior controls in this case are bang-bang.

#### Theorem 17.

*If α* = 0, *there are no singular arcs for the optimal time problem presented in Section 4. Thus, the interior control structure is bang-bang*.

Outside of the singular arc 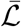, the control structure is completely determined by (66) and (84). The precise result, utilized later for optimal synthesis presented in Section 7, is stated in the following theorem. We first introduce a convenient (and standard) notation. Let finite words on *X* and *Y* denote the concatenation of controls corresponding to vector fields *X*(*u* ≡ 0) and *Y* (*u* ≡ *M*), respectively. The order of application is read left-to-right, and an arc appearing in a word may not actually be applied (e.g. *XY* denotes an *X* arc followed by a *Y* arc or a *Y* arc alone).

#### Theorem 18.

*Consider an extremal lift* Γ = ((*x, u*), λ). *Trajectories x remaining entirely in* 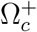 *or* 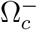 *can have at most one switch point. Furthermore, if* 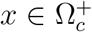, *then the corresponding control is of the form YX. Similarly*, 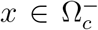 *implies that u = XY. Hence multiple switch points must occur across the singular arc* 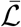.

*Proof*. If *τ* is a switching time, so that Φ(*τ*) = 0, equation (84) allows us to calculate 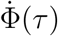 as

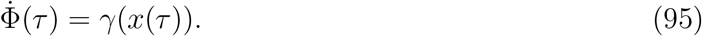

Thus, in 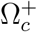 where *γ* > 0, 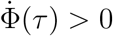, and hence Φ must increase through *τ*. The expression for the control (66) then implies that a transition from a *Y*-arc to an *X*-arc occurs at *τ* (i.e. a *YX* arc). Furthermore, another switching time cannot occur unless *x* leaves 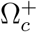, since otherwise there would exist 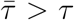 such that 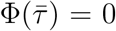, 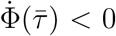 which is impossible in 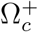. Similarly, only *XY*-arcs are possible in 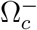.

The structure implies by Theorem 18 is illustrated in Figure 3. Note that inside the sets 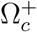, 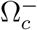, and 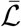, extremal lifts are precisely characterized. Furthermore, the results of Section 6.1 (and particularly equation (60)) yield the characterization only the boundary *N*. What remains is then to determine the synthesis of these controls to the entire domain Ω_*c*_, as well as to determine the order local optimality of the singular arc 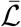. The latter is addressed in the following section.

### 6.4 Optimality of Singular Arcs

We begin by proving that the singular arc is extremal, i.e. that it satisfies the necessary conditions presented in Section 6.2 (note that it is interior by assumption). This is intuitively clear from Figure 3, since *X* and *Y* point to opposite sides along 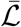 by the defintion 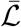.

#### Theorem 19.

*The line segment* 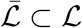 *is a singular arc*.

*Proof*. We find an expression for *u* = *u*(*x*) such that the vector *f*(*x*) + *u*(*x*)*g*(*x*) is tangent to 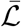 at *x*, i.e. we find the unique solution to

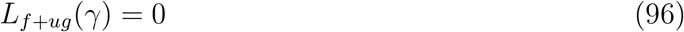

Note that we can invert (91):

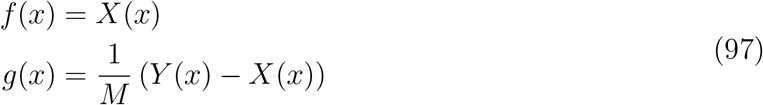

so that 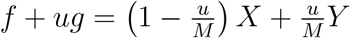. Thus,

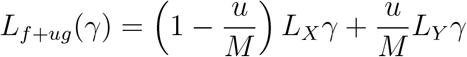

Setting the above equal to zero, and solving for *u* = *u*(*x*) yields

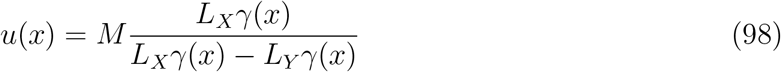

As *L_Xγ_* < 0 and *L_Yγ_* > 0 on 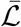 by Proposition 15, we see that 0 < *u*(*x*) < *M*. We must also verify that the associated controlled trajectory (98) is extremal by constructing a corresponding lift. Suppose that *x*(*t*) solves

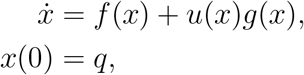

for 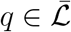. Let *ϕ* ∈ (ℝ^2^)^*^ such that

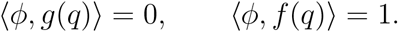

Let λ(*t*) solve the corresponding adjoint equation (65) with initial condition λ(0) = *ϕ*. Then the extremal lift Γ = ((*x,u*),λ) is singular if Φ(*t*) = 〈λ(*t*), *g*(*x*(*t*))〉 ≡ 0. By construction of *u*(*x*), the trajectory remains on 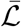 on some interval containing zero, and we can compute 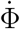 as (using (75))

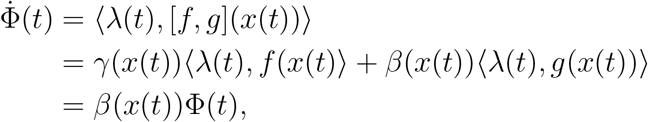

Note that we have used (84) and the fact that *γ* = 0 by our choice of *u*. Since Φ(0) = 0 by hypothesis, this implies that Φ(*t*) ≡ 0, as desired.

The above then verifies that 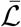 is a singular arc. Note that an explicit expression for *u* = *u*(*x*) was given in (93), which can be shown to be equivalent to (98).

Having shown that the singular arc 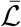 is extremal, we now investigate whether it is locally optimal for our time-optimization problem. The singular arc is of intrinsic order *k* if the first 2*k* – 1 derivatives of the switching function are independent of *u* and vanish identically on an interval *I*, while the 2*k*^th^ derivative has a linear factor of *u*. We can compute (this is standard for control-affine systems (5)) that

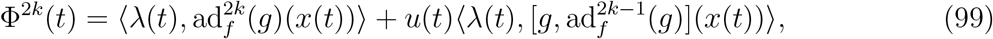

where ad_*Z*_ is the adjoint endomorphism for a fixed vector field *Z*:

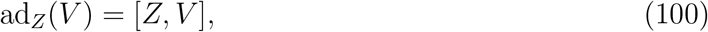

and powers of this operator are defined as composition. Fix an extremal lift Γ = ((*x, u*), λ) of a singular arc of order *k*. The Generalized Legendre-Clebsch condition (also known as the Kelley condition) [13] states that a necessary condition for Γ to satisfy a minimization problem with corresponding Hamiltonian *H* is that

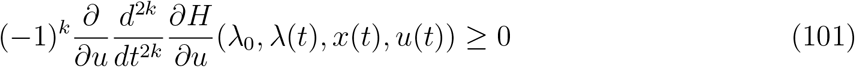

along the arc. Note that 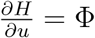, so that the above is simply the *u* coefficient of the 2*k*th time derivative of the switching function (multiplied by (–1)^*k*^). The order of the arc, as well as the Legendre-Clebsch condition, are addressed in Theorem 20.

#### Theorem 20.

*The singular control is of order one. Furthermore, for all times t such that* 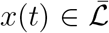, 〈λ(*t*), [*g*, [*f, g*]](*x*(*t*))〉 > 0. *Thus, the Legendre-Clebsch condition is violated, and the singular arc* 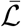 *is not optimal*.

*Proof*. Along singular arcs we must have Φ(*t*), 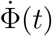, 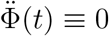, and we can compute these derivatives using iterated Lie brackets as follows:

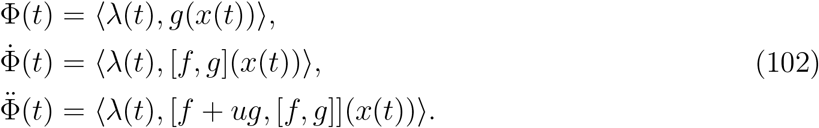

The final of the above in (102) can be simplified as

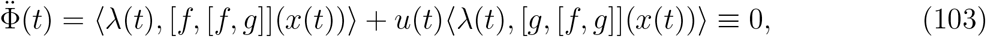

which is precisely (99) for *k* = 1. Order one is then equivalent to being able to solve this equation for *u*(*t*). Thus, 〈λ(*t*), [*g*, [*f,g*]](*x*(*t*))〉 > 0 will imply that the arc is singular of order one. We directly compute 〈λ(*t*), [*g*, [*f, g*]](*x*(*t*))〉 = 〈λ(*t*), [*g*, ad_*f*_(*g*)](*x*(*t*))〉. Using equation (75) and recalling properties of the singular arc (*γ* = 0 and the remaining relations in (102), as well as basic “product rule” properties of the Lie bracket), we can show that

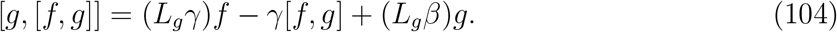

Recall that for an extremal lift along the arc 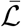,

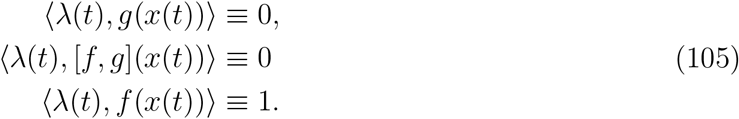

The first two of the above follow from Φ, 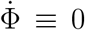, and the third is a consequence of *H* ≡ 0 (see (63)). Equations (104) and (105) together imply that

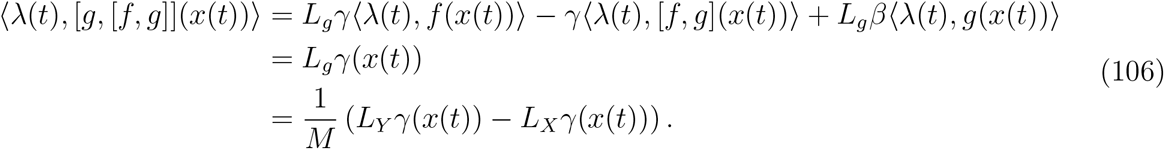

The last equality follows from the representation in (97). As *L_Yγ_* > 0 and *L_Xγ_* < 0 along 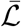 (Proposition 15), 〈λ(*t*), [*g*, [*f, g*]](*x*(*t*))〉 > 0, as desired. Furthermore,

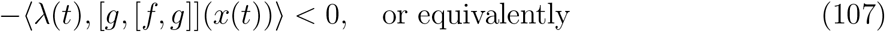

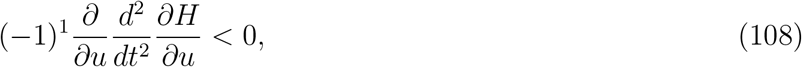

showing that (101) is violated (substituting *k* = 1). Thus, 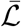 is not optimal.

Theorem 20 then implies that the singular arc is suboptimal, i.e. that 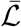 is “fast”with respect to the dynamics. In fact, comparing time along the trajectories can be computed explicitly using the “clock form,” a one-form on Ω. As one-forms correspond to linear functionals on the tangent space, and *f* and *g* are linearly independent, there exists a unique *ω* ∈ (*T*Ω)^⋁^ such that

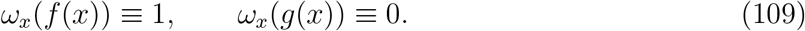

In fact, we compute it explicitly:

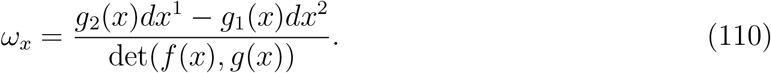

Then, along any controlled trajectory (*x, u*) defined on [*t*_0_, *t*_1_], the integral of *ω* computes the time *t*_1_ – *t*_0_:

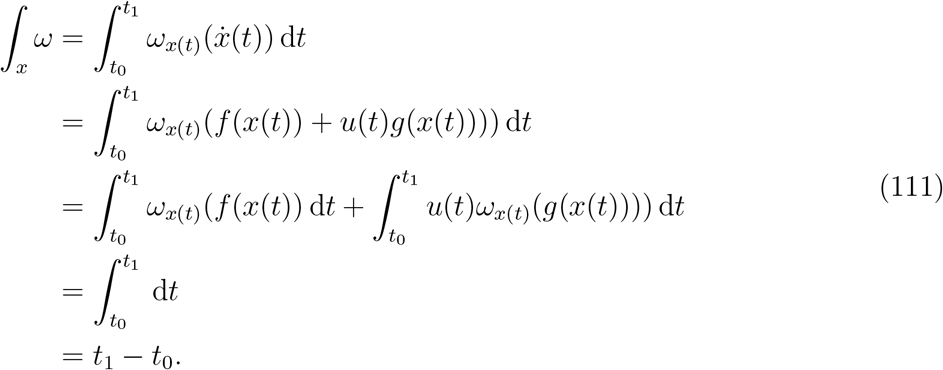

We can then use *ω* and Stokes’ Theorem to compare bang-bang trajectories with those on the singular arc. See Figure 4 below for a visualization of a singular trajectory connecting *q*_1_, 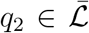 and the corresponding unique *XY* trajectory connecting these points in 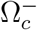 (note that uniqueness is guaranteed as long as *q*_1_ and *q*_2_ are sufficiently close).

**Figure 4:**
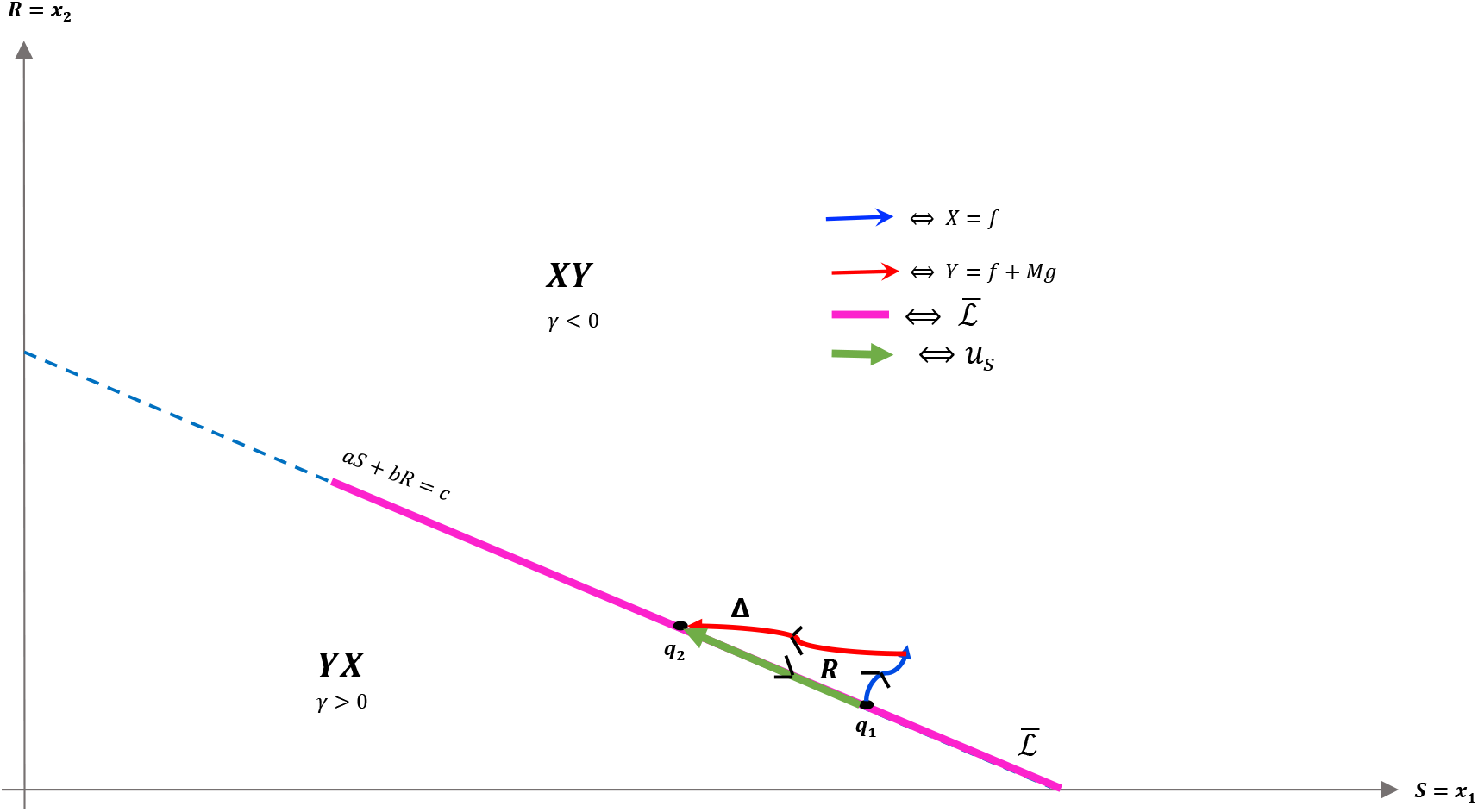
Both *XY* and singular trajectories taking *q*_1_ to *q*_2_.

Let *t_S_* denote the time spent along the singular arc, *t_X_* the time spent along the *X* arc, and *t_Y_* the time spent along the *y* arc. Denote by Δ the curve traversing the *X* and *y* arcs positively, and the singular arc negatively, and *R* its interior. As *X* and *Y* are positively oriented (they have the same orientation as *f* and *g*), Stokes’ Theorem yields

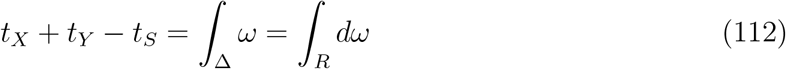

Taking the exterior derivative yields the two-form *dω* (see Chapter 2 of [13]):

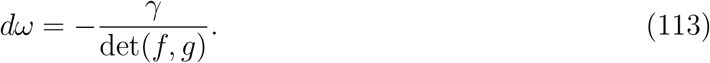

As the determinant is everywhere positive (see the proof of Proposition 12), and *R* lies entirely in *γ* < 0, the integral on the right-hand side of (112) is positive, so that we have

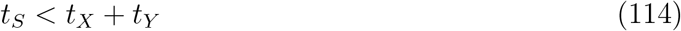

Thus, time taken along the singular arc is shorter than that along the *XY* trajectory, implying that the singular arc is locally suboptimal for our problem (recall that we want to maximize time). Since local optimality is necessary for global optimality, trajectories should never remain on the singular arc for a measurable set of time points. This reaffirms the results of Theorem 20. A completely analogous statement holds for *YX* trajectories in the region *γ* > 0. We can also demonstrate, utilizing the same techniques, that increasing the number of switchings at the singular arc speeds up the trajectory; see Figure 5. This again reinforces Theorem 20, and implies that trajectories should avoid the singular arc to maximize the time spent in Ω_*c*_.

**Figure 5:**
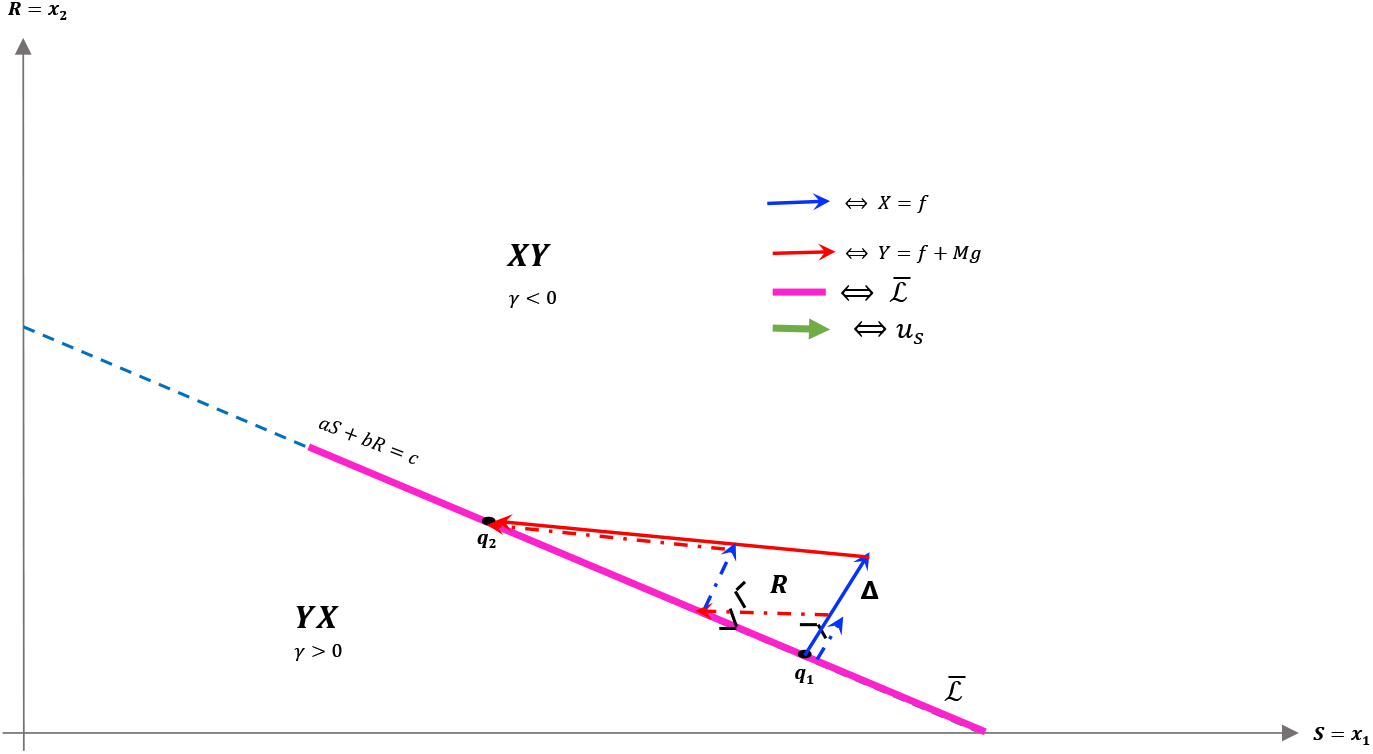
*XY* (solid) and *XY XY* (dashed) tra jectories taking *q*_1_ to *q*_2_ in the region *γ* > 0. The time difference between the two trajectories can again be related to the surface integral in the region *R*, where *γ* < 0. The *XY* trajectory can then be seen to be slower in comparison.

## 7 Characterization of Optimal Control

The results of Section 6.1, 6.2, 6.3, and 6.4 may now be combined to synthesize the optimal control introduced in Section 4. First we provide a lemma which shows that arcs of the form *Yu_p_* are not possible.

### Lemma 21.

*If the optimal control has a switch point in* 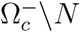, *then the optimal control does not slide (i. e. there is not t*_1_, *t*_2_ *with* 0 < *t*_1_ < *t*_2_ < *t_c_ such that* 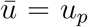 *on the open interval* (*t*_1_, *t*_2_)).

*Proof*. This result follows from the fact that in 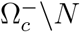 the optimal control can switch one time, and only from *u* = 0 to *u* = *M*. Once *u* = *M*, the only way the optimal trajectory would reach *N*, so it may be able to slide, is if *L_Y_V*(*x*) > 0 for 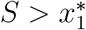.

Given *δ* > 0 small enough, suppouse there is a trajectory *x*(*t*) = (*S*(*t*), *R*(*t*)) and a time *t_δ_* such that

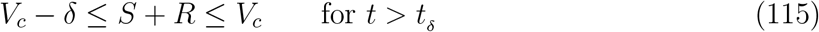

and *L_Y_V*(*x*) > 0. From equation (115) and the definition of *L_Y_V*(*x*) it satisfies

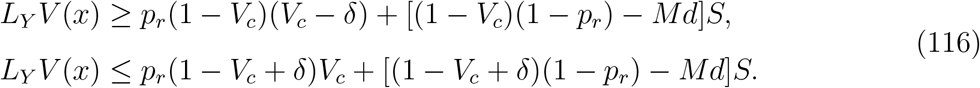

Notice that the second inequality on equation (116) implies that

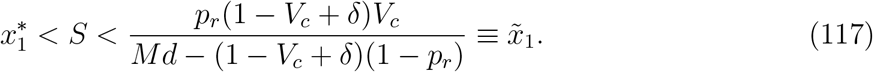

On the other hand, the continuity of the state variables and the assumption that *L_Y_V*(*x*) > 0 for all *δ* > 0 small enough implies that

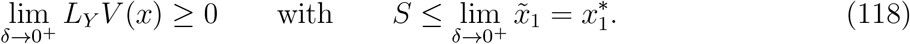

Thus, on the *Y* arc, the *x*(*t*) trajectory will reach *N* only if 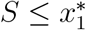. According to Proposition 7, *u_p_* is unfeasible on this region. Therefore, the trajectory does not slide on [0, *t_c_*].

### Theorem 22.

*For any α* ≥ 0, *the optimal control to maximize the time to reach a critical time is a concatenation of bang-bang and path-constraint controls. In fact, the general control structure takes the form*

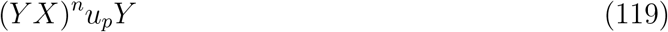

*where* (*YX*)^*n*^:= (*YX*)^*n*−1^*YX for n* ∈ ℕ, *and the order should be interpreted as left to right. Here up is defined in* (60).

*Proof*. Formula (119) is simply a combination of the results presented previously. Note that singular arcs are never (locally) optimal, and hence do not appear in the equation. We also observe that *X* arcs are not admissible once the boundary *N* has been obtained, as an *X* arc always increases *V*. A *Y* arc may bring the trajectory back into int(Ω_*c*_), but a *YX* trajectory is no longer admissible, as the switching structure in Ω_−_ is *XY* (Theorem 18). Note that Lemma 21 implies that a *Y* arc is not necessary after the *n* interior switches (*YX*)^*n*^.

The only aspect that remains is to show that once *N* is reached, the only possible trajectories are either *u_p_* given by (60) or *Y*, with at most one switching occurring between the two. That is, a local arc of the form *u_p_Yu_p_* is either sub-optimal or non-feasible (equivalently, outside of the control set *U*). Suppose that such an arc is feasible, i.e. that for all such points in phase space, 0 ≤ *u_p_* ≤ *M* (recall that up is defined via feedback in (60)). Denote by *τ*_1_ and *τ*_2_ the times at which the switch onto and off of *Y* occurs, respectively. Since up decreases with *S*, feasibility implies that *u_p_*(*τ*_1_), *u_p_*(*τ*_2_) ≤ *M* and furthermore that *u_p_*(*t*) ≤ *M* for all *t* ∈ [*τ*_1_, *τ*_2_]. Thus, we can consider the alternate feasible trajectory which remains on *N* between the points (*S*(*τ*_1_), *R*(*τ*_1_)) and (*S*(*τ*_2_), *R*(*τ*_2_)); see Figure 6 for an illustration. Call *τ* the time for such a trajectory. Then, using the clock-form *ω* and the positively-oriented curve Δ which follows *N* first and *Y* (in the reverse direction) second, we obtain similarly to (112),

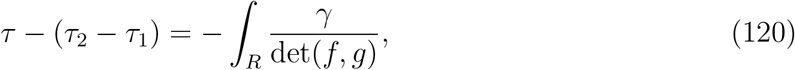

where *R*:= int(Δ). Recalling that *γ* < 0 in *R* (see Figure 3), the previous equation implies that

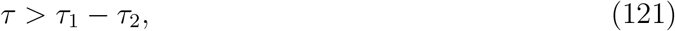

i.e. a longer amount of time is achieved by remaining on the boundary *N*. Hence the arc *u_p_Yu_p_* is sub-optimal if it is feasible, as desired.

The previous argument has one subtle aspect, as we used results from the Maximum Principle on the boundary set *N*, where technically it does not apply. However, the above still remains true, since we may approximate the boundary line *V* = *V_c_* with a curve interior to Ω_*c*_ which remains feasible. By continuity, the time along such a curve can be made arbitrarily close to *τ*, and hence is still greater than *τ*_2_ – *τ*_1_, implying that *u_p_Yu_p_* is sub-optimal.

**Figure 6:**
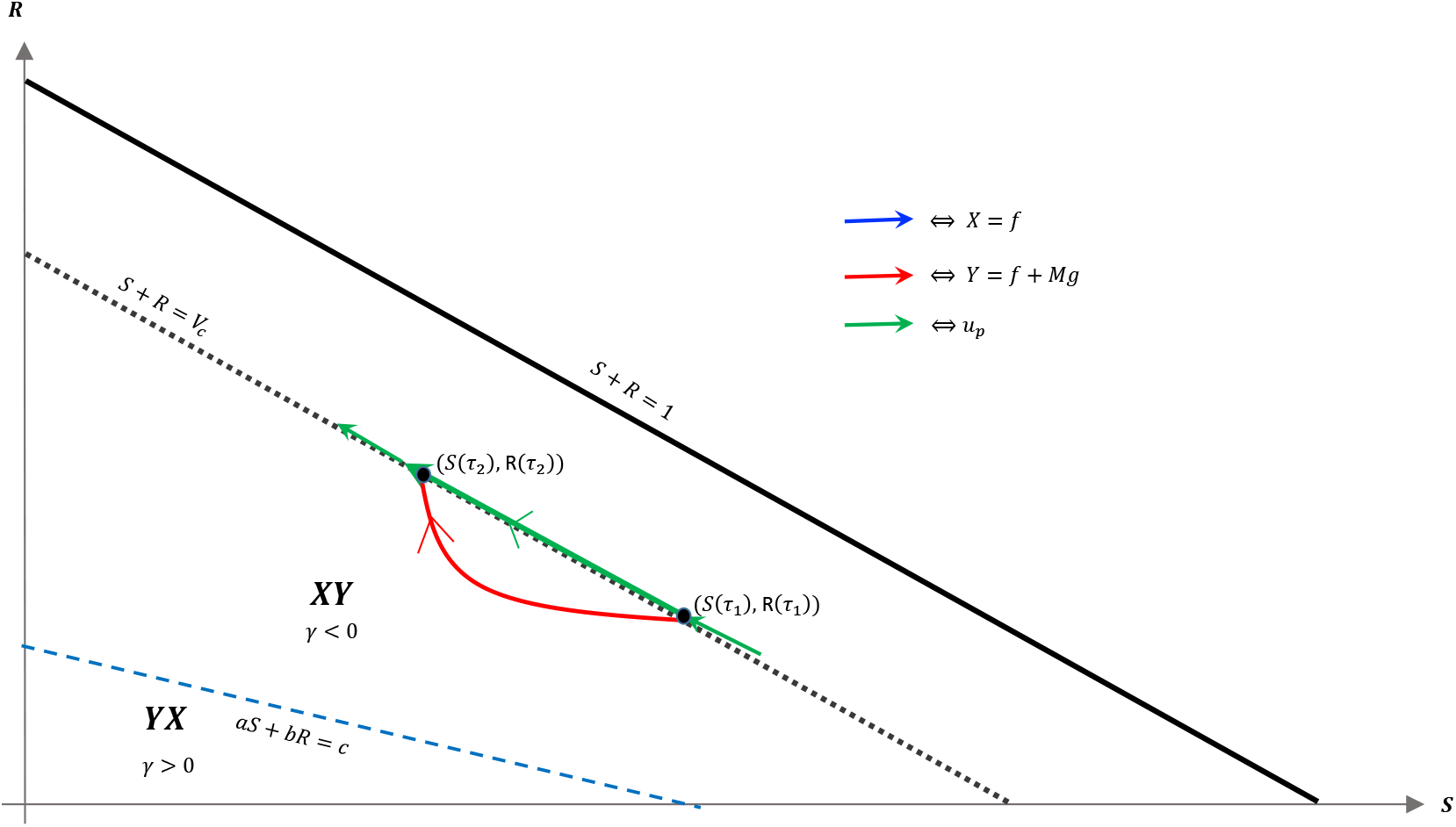
Comparison of *u_p_Yu_p_* arc and an arc that remains on *N* (hence *u* ≡ *u_p_*) between the points (*S*(*τ*_1_), *R*(*τ*_1_)) and (*S*(*τ*_2_), *R*(*τ*_2_)), assuming that up remains feasible (that is, *u_p_* ∈ [0, *M*]). Note that *γ* < 0 in the area of interest, and that a switching of a *Y* to an *X* arc is prohibited via the Maximum Principle. Thus, the only possibility is the curve illustrated, which leaves the boundary *N* for a *Y* arc before *u_p_* becomes infeasible.

### Proof option 2 to Theorem 22

Formula (119) is simply a combination of the results presented previously. Note that singular arcs are never (locally) optimal, and hence does not appear in the equation. We also observe that *X* arcs are not admissible once the boundary *N* has been obtained, as an *X* arc always increases *V*. A *Y* arc may bring the trajectory back into int(Ω_*c*_), but a *YX* trajectory is no longer admissible, as the switching structure in Ω_−_ is *XY* (Theorem 18). Thus, once *N* is reached, the only possible trajectories are either *u_p_* given by (60) or *Y*. Observe that no more than one switch point exists, otherwise, a local arc will have the form: *u_p_Yu_p_* or *Yu_p_Y*, which implies the existence of a switch point from *Y* to *u_p_*, contradicting results shown on the proof of Lemma 21.

Note that in Theorem 22, the switchings must occur across the singular arc 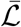, if it exists (recall that it is not admissible if *α* = 0). The control up is determined along the boundary of Ω_*c*_, and provides the synthesis between exterior and boundary controls.

## 8 Numerical Results

In this section, we provide numerical examples of the analytical results obtained in previous sections. All figures in this section were obtained using the GPOPS-II Matlab software. Recall the label *x*_1_ and *x*_2_ for the sensitive and resistant cells, respectively. In all experiments we utilize the parameters and initial values given in Table 1 below, unless stated otherwise.

**Table 1:**
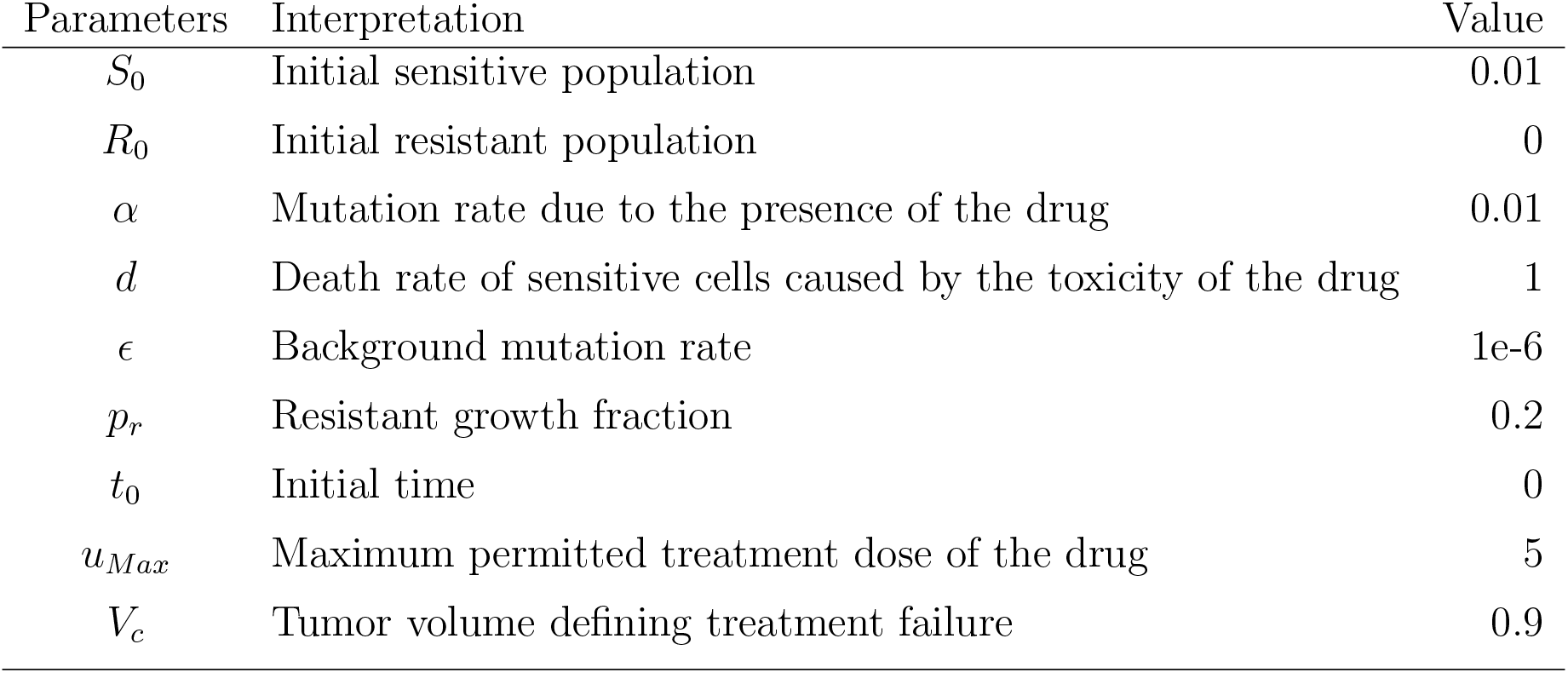
Parameter values and initial conditions used in most of the figures of section 8 unless otherwise stated in the caption of the figure.

Theorem 22 characterizes the qualitative form of the optimal control:

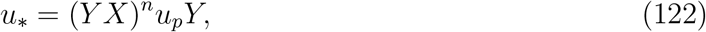

where *n* is the number of interior switches, *u_p_* the sliding control (60), and *X* and *Y* denote the boundary controls *u* = 0 and *u* = *M*, respectively. We begin by computing sample controls; see Figures 7 and 9. Note that the optimal control in Figure 7(b) takes the form *YXu_p_Y*, while that of Figure 9(b) is an upper corner control *Y*. The phase plane dynamics corresponding to Figure 7 is also provided in Figure 8. In both cases the cytotoxic parameter was fixed at *d* = 0.05, while the induced rate of resistance *α* varies between *α* = 0.005 in Figure 7 and *α* = 0.1 in Figure 9. Note that for the smaller value of *α* (Figure 7), a longer period of treatment success is observed, as the time to treatment failure is approximately 70 time units; compare this with *t_c_* = 24.2 in Figure 9. This result may be expected, as the less mutagenic drug is able to be more effective treatment when optimally applied.

**Figure 7:**
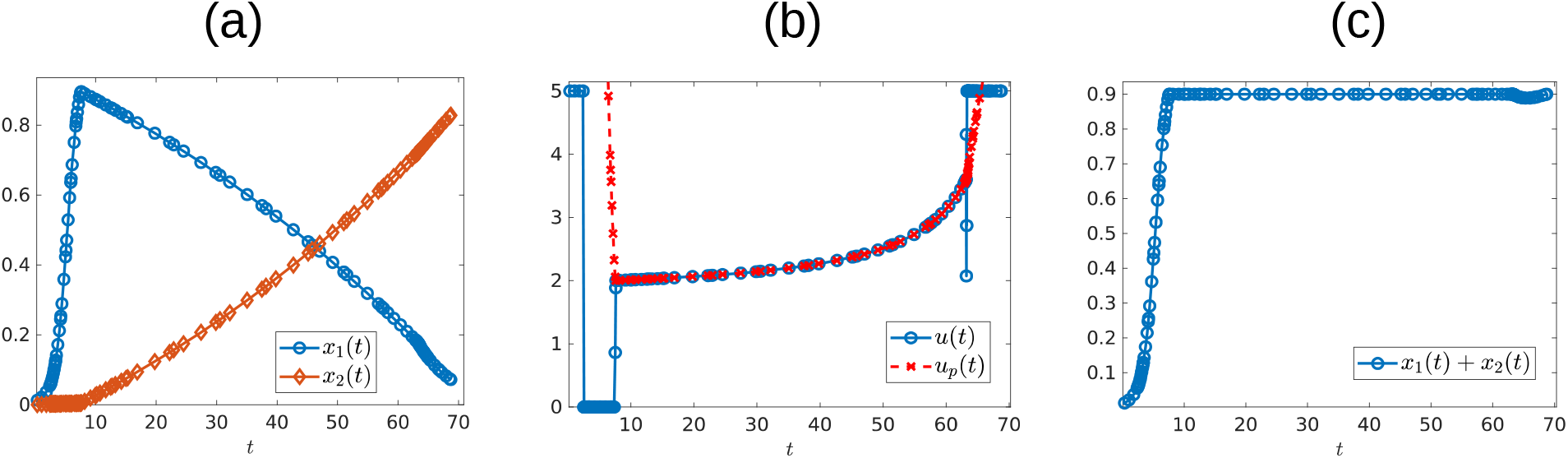
Numerical solution of optimal control problem with *d* = 0.05, *α* = 0.005, and the remainder of parameters as in Table 1. (a) Sensitive (*x*_1_) and resistant (*x*_2_) temporal dynamics. (b) Control structure of form *YXu_p_Y*. (c) Volume dynamics. Note that the trajectory remains on the line *V* = *V_c_* for most times, with corresponding control *u* = *u_p_*.

**Figure 8:**
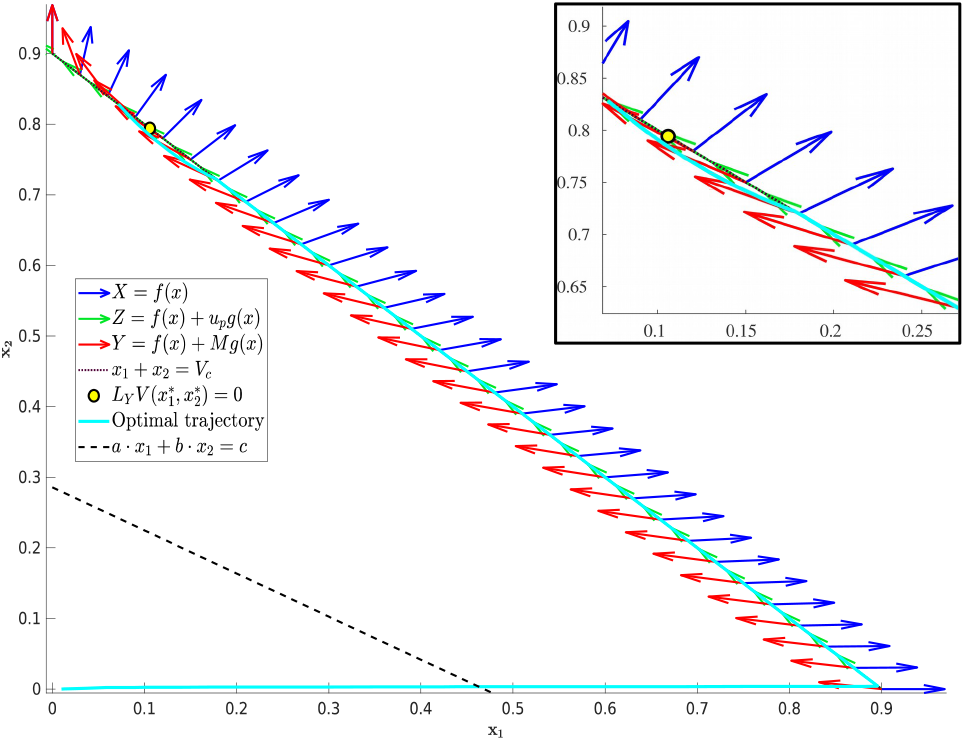
Phase plane corresponding to Figure 7. Trajectory which optimal control is of the form *YXu_p_Y* with parameter values as in table 1 except for *α* = 0.005 and *d* = 0.05. The yellow dot in the figure represents the 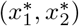 point at which *Y*(*x*) is tangent to the sliding surface. Here, 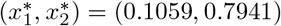. As proved in theorem 6, for points on the line *N* the volume will decrease when it moves on the *Y*(*x*) direction if *x*_1_ > 0.1059 and will increase for *x*_1_ < 0.1059.

**Figure 9:**
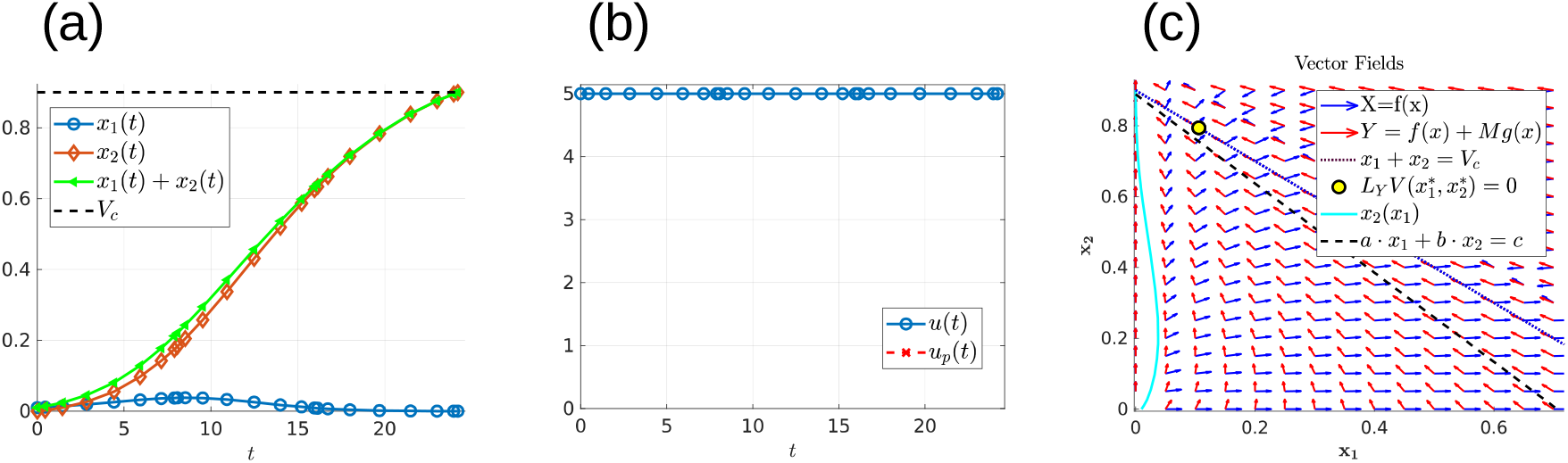
Numerical solution of optimal control problem with *d* = 0.05, *α* = 0.1, and the remainder of parameters as in Table 1. (a) Sensitive (*x*_1_), resistant (*x*_2_), and volume (*x*_1_ + *x*_2_) temporal dynamics. (b) Control structure of form *Y*, i.e. an entirely upper corner control. (c) Phase plane dynamics, plotted with relevant vector fields.

The generality of the previous statement is investigated in Table 2 and Figures 10 and 11. The computed optimal times *t_c_* suggest that when the cytotoxicity of the drug (d) is small, higher induction rates (*α*) actually increase treatment efficacy. For example, for *d* = 0.001 success is maximized when *α* takes larger values (Figure 11(a)). However, the situation appears to be reversed when we consider larger values of *d*; see for example the row *d* = 0.5 in Table 2, and the corresponding purple curve in Figure 11(b). Figure 11(b) actually provides the critical time as a function of *α* for multiple cytotoxicities *d*, and we observe the qualitative differences as *d* increases. Examining Figure 10 and Table 2 also suggests that as *d* increases, the feedback control up becomes optimal on an interval [*t*_1_, *t*_2_] with 0 < *t*_1_ < *t*_2_ < *t_c_*.

**Figure 10:**
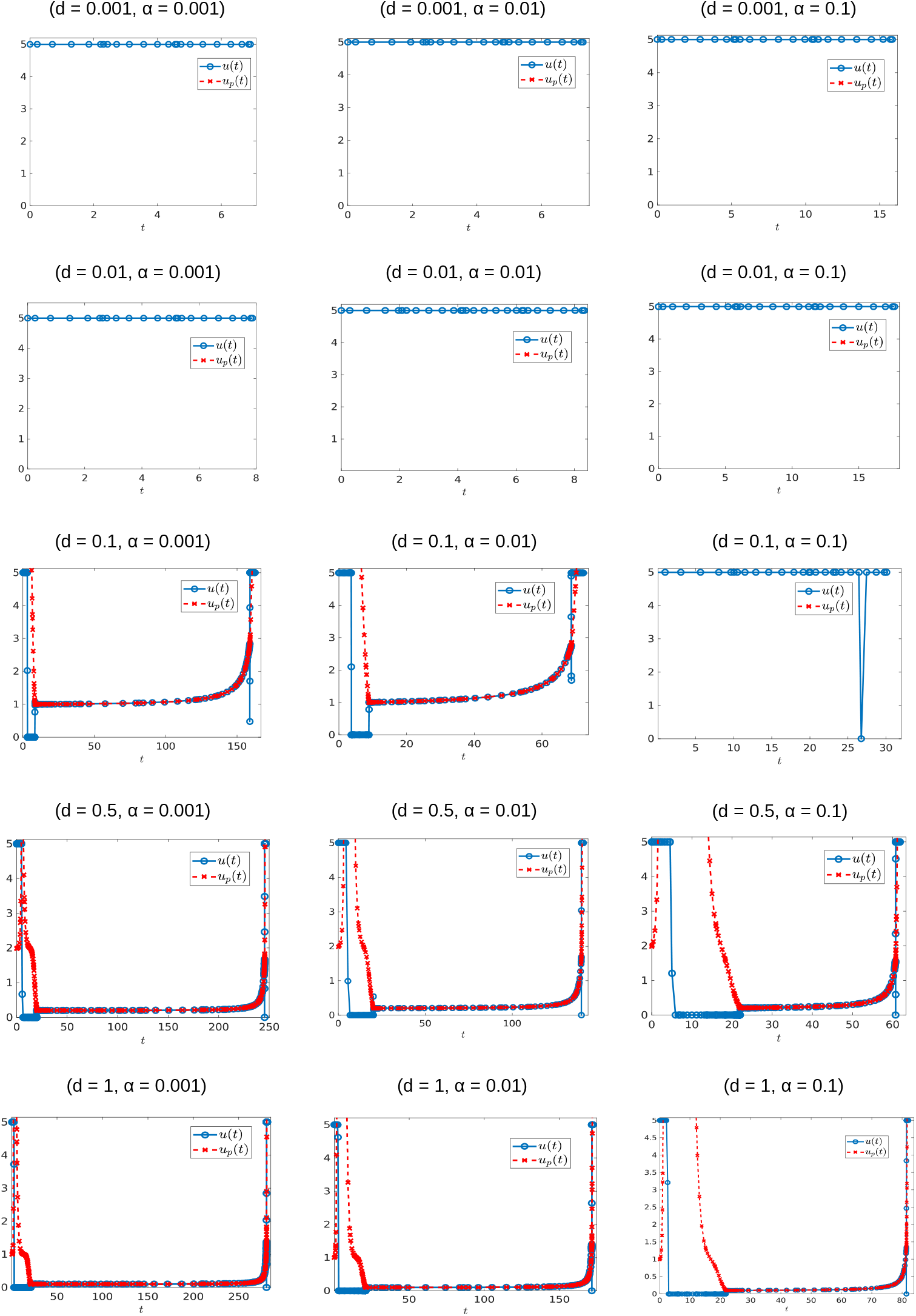
Optimal control structures for different *α* and *d* values. The blue curve is the optmimal control, the red curve is the feedback control that may or may not be optimal or even feasible.

**Figure 11:**
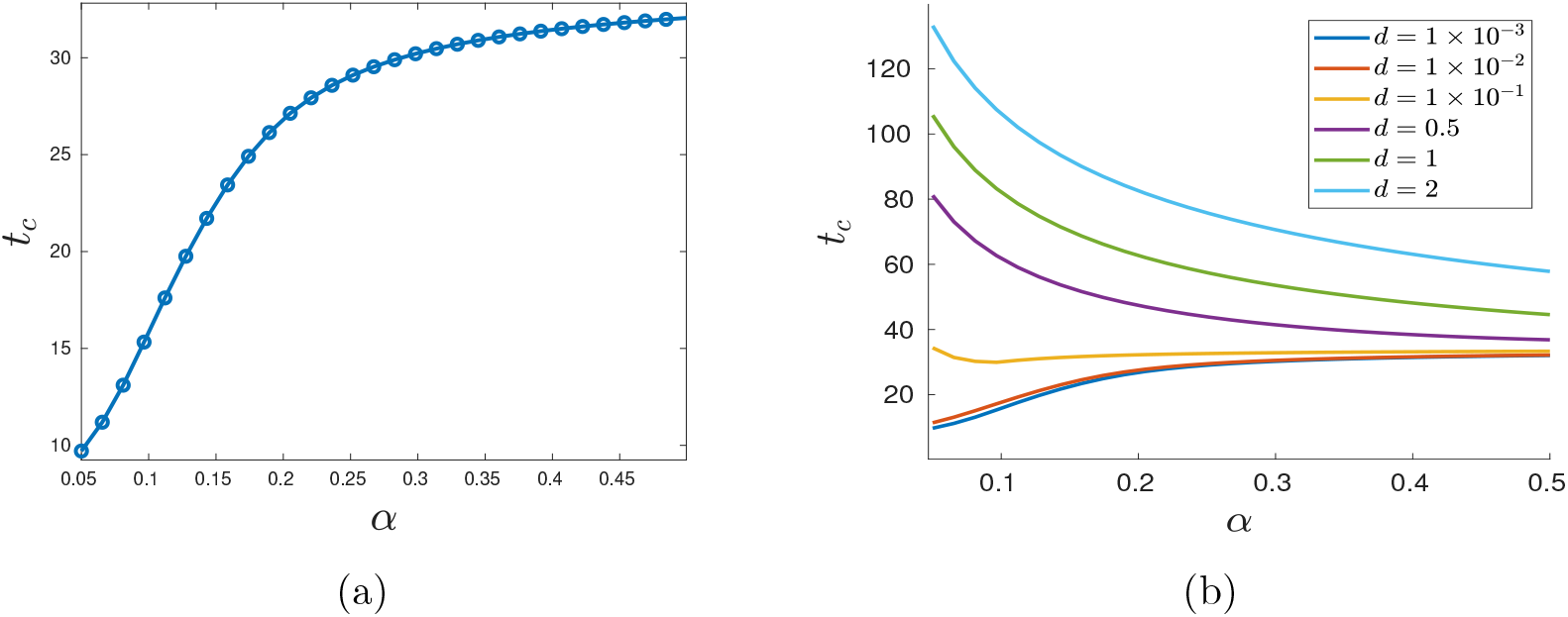
Variation in *t_c_* as a function of *α*. (a) Treatment success time *t_c_* for *d* = 0.001 with varying *α* values. (b) Functional dependence of *t_c_* on *α* for different *d* parameters. Note that for small *d, t_c_* increases as a function of *α*, but that this trend is reversed if *d* is further increased.

**Table 2:**
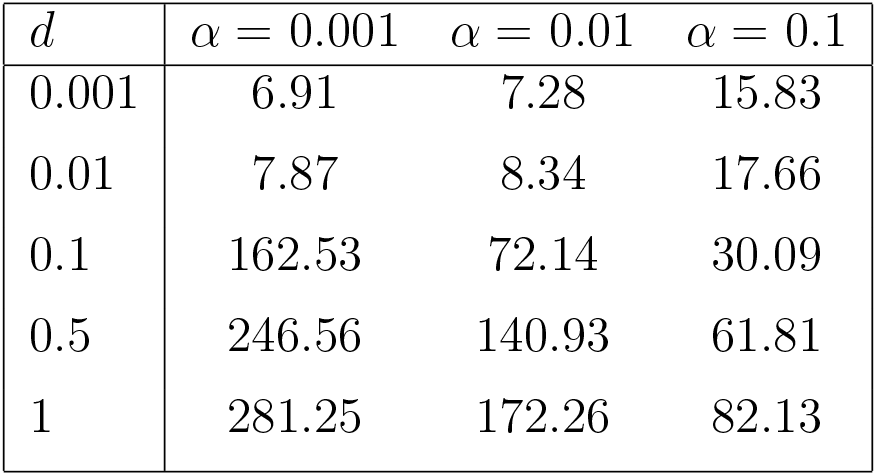
Optimal maximal end time *t_c_* for each of the computed controls appearing in Figure 10.

We similarly investigated the control structure and treatment outcome as a function of *d* for a fixed *α*; these results are presented in Figures 12 and 13. Here *α* = 0.005 is fixed and *d* is varied on the interval [0.001, 0.1]. Figure 12 presents three of these controls; although none of the controls is of the form *YXY*, the figure suggests that there may exist a *d*^*^ ∈ (0.02062, 0.0207959) where the solution trajectory may intersect the boundary line *N* only at one point and subsequently switch into a *Y* arc, thus providing the existence of a *YXY* control. Figure 13 suggests that increasing *d* for a fixed *α* increases the overall effectiveness of the treatment for all values of *α*, and that decreasing the induction rate *α* allows for longer tumor control. However, for small values of *d*, increasing *α* may provide a better treatment regime (see, for example, the yellow and purple curves in Figure 13).

**Figure 12:**
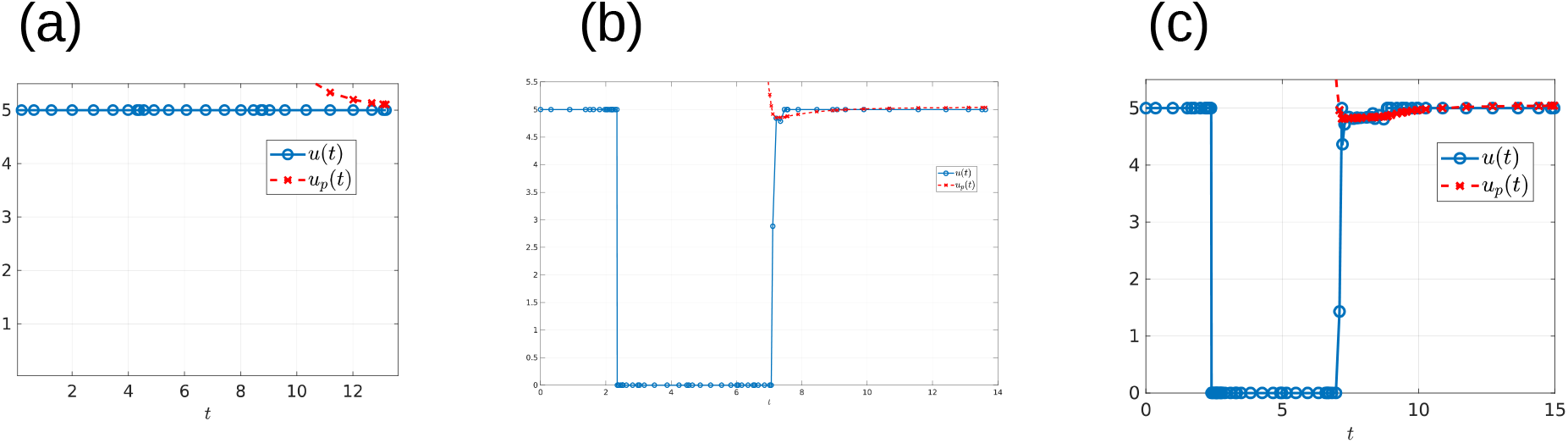
Computed optimal controls for *α* = 0.005 and (a) *d* = 0.0206, (b) *d* = 0.020624489795918 and (c) *d* = 0.207959. Note that the control in (a) takes the form *Y*, while that in (b) and (c) is of the form *YXu_p_*.

**Figure 13:**
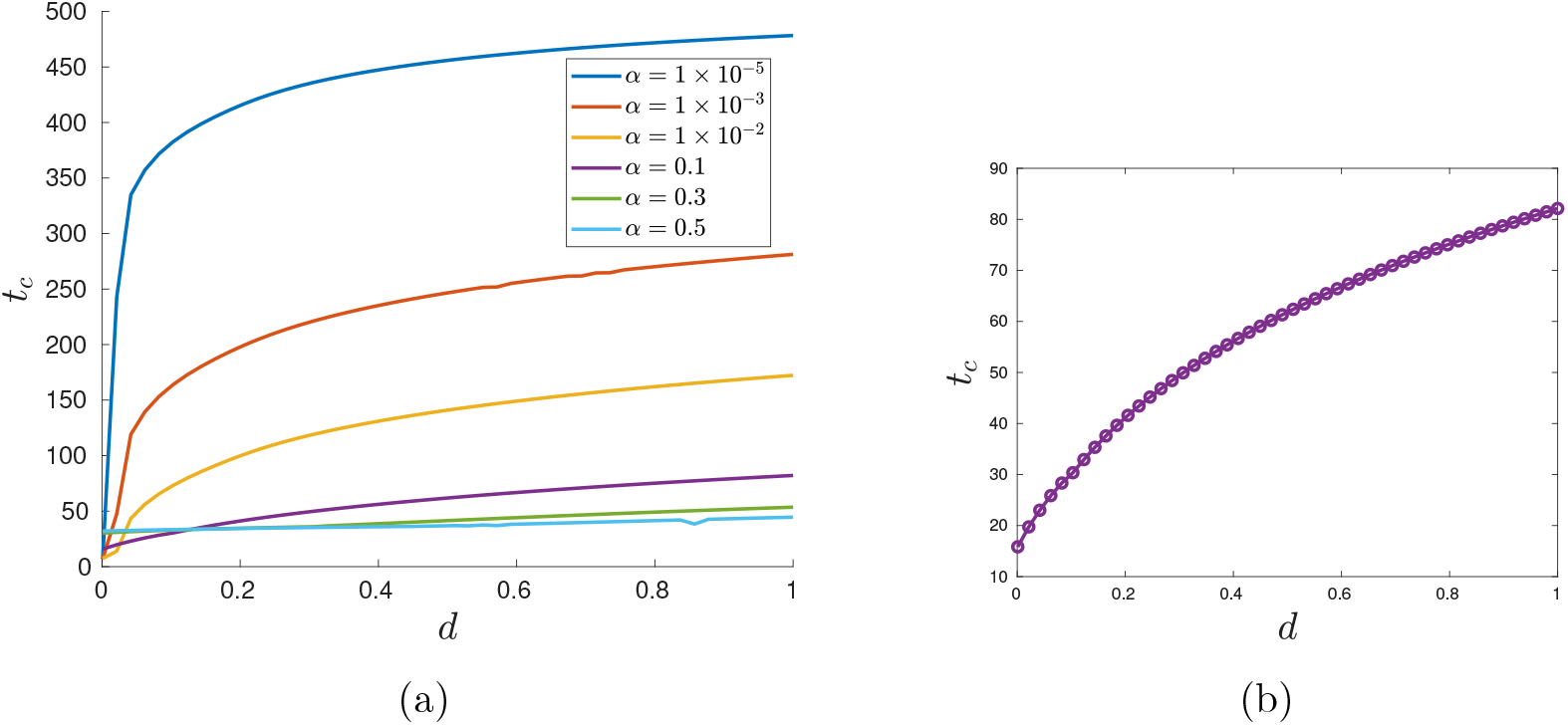
Variation in *t_c_* as a function of *d*. Parameter *α* is fixed at *α* = 0.005 (a) *t_c_* response for varying *d* values. Note that treatment efficacy generally increases with increasing *d*. (b) *α* = 0.1.

Lastly, to verify the performance of the numerical software, we regularize the control problem to include a quadratic cost with respect to the control. More precisely, we consider the perturbed performance index

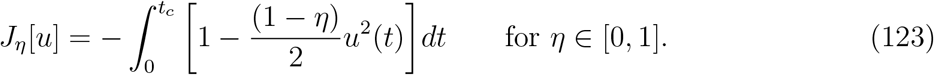

Notice that equation (123) represents a family of performance indexes parameterized by *η*. Furthermore, the original performance index corresponds to *η* = 1. We investigate the corresponding control structure in the limit *η* → 1. For *η* ≠ 1, the optimal control problem is regular and solvers such as GPOPS-II, SNOPT or a single shooting method should provide accurate solutions. We begin with the solution of the regular problem (say *η* = 0) and let *η* → 1 to attain the solution of the original problem formulated in Section 4. Figure 14 provides the computed controls as perturbation constant *η* takes the values 0, 0.5, 0.7, 0.9, 0.95, 0.999, 0.99999 and 1. For each case we obtained different relative errors with 4.0338 × 10^™7^ the largest relative error (on case *η* = 0), the remaining values of *η* have smaller relative errors. It is clear, from Figures 14(case *η* = 0.95), 14(case *η* = 0.999) and 14(case *η* = 0.99999) that as *η* → 1 the control approaches the solution to the original problem (compare to Figure 7).

**Figure 14:**
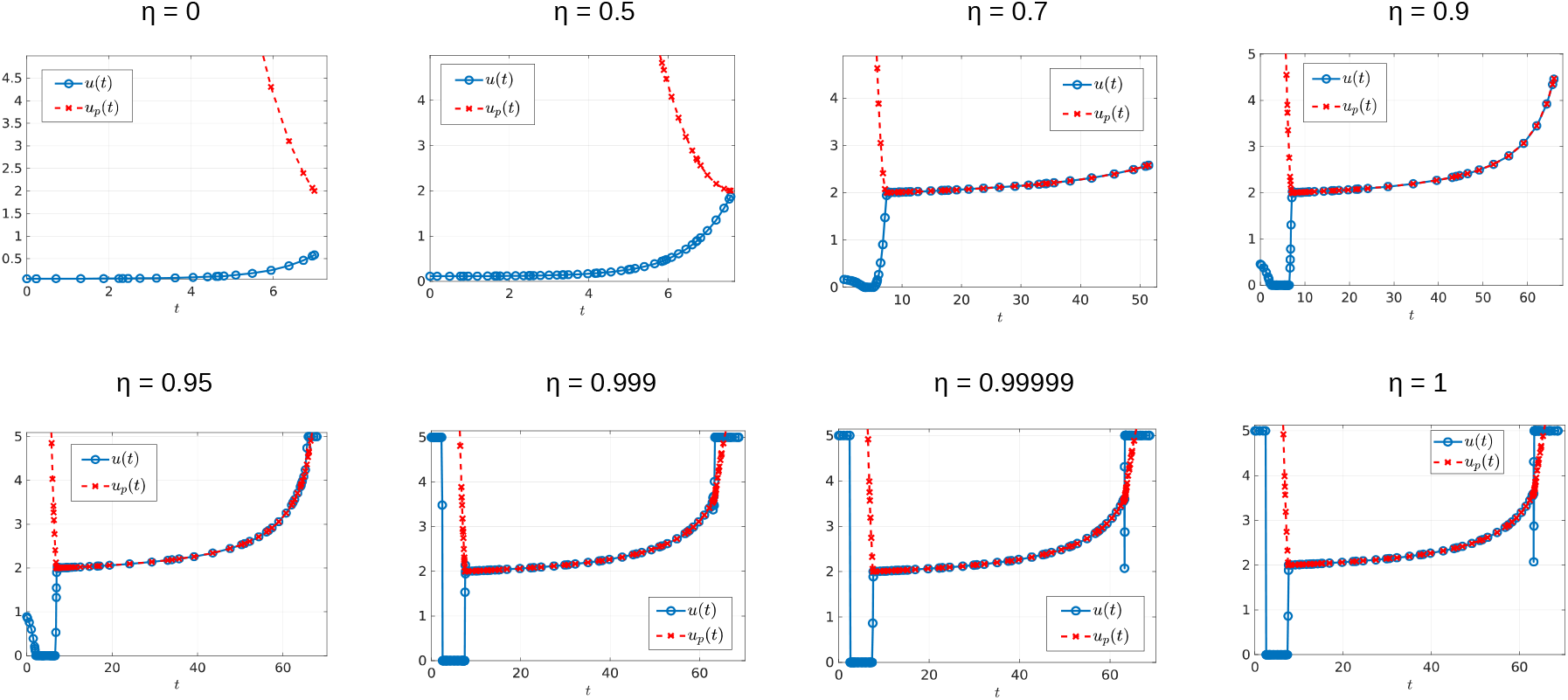
Different perturbed controls for *α* = 0.005 and *d* = 0.05. Here, from Figures (a) to (h), the value of *η* is 0, 0.5, 0.7, 0.9, 0.95, 0.999, 0.99999 and 1, respectively. The maximum relative error is of 4.0338 × 10^−7^ for figure *η* = 0, the remaining figures have a maximum relative error of 5.5727 × 10^−7^ or smaller.

## 9 Conclusions

In this work, we have provided the mathematical details presented in [10]. That is, we have formally applied optimal control theory techniques to understand treatment strategies related to a model of induced drug resistance in cancer chemotherapy introduced in [11]. We have shown that a drug’s level of resistance induction is identifiable, thus allowing for the possibility of designing therapies based on individual patient-drug interactions. An optimal control problem is then presented which maximizes a specific treatments therapy window. Existence results are established, and a formal analysis of the optimal control structure is performed utilizing the Pontryagin Maximum Principle and differential-geometric techniques. Optimal treatment strategies are realized as a combination of bang-bang and path-constrained arcs, and singular controls are proved to be sub-optimal. Numerical results are presented which verify our theoretical results, and demonstrate interesting and non-intuitive treatment strategies.

The numerical results of Section 8 provide interesting behavior of the optimal control as a function of parameter values. For example, the behavior of the objective *t_c_* depends on the value of the drug-kill rate *d*: for large *d, t_c_* increases with induction-rate *α*, while this behavior is reversed for small *d* (Figure 11). Also, our simulations suggest that it is optimal to apply the maximal dosage *M* subsequent to sliding along the boundary *V* = *V_c_* (e.g. Figure 8), prior to treatment failure. Further analysis is required to understand this phenomenon, and its implications for clinical scenarios.

Other questions remain open for future work:

⋄ For controls where the trajectory remains on the boundary *V* = *V_c_* (*u_p_*), the feedback control is optimal during a time interval [*t*_1_, *t*_2_] with 0 ≤ *t*_1_ < *t*_2_ < *t_c_*. It remains to understand the point of entry (*x*_1_(*t*_1_), *x*_2_(*t*_1_)) and exit (*x*_1_(*t*_2_), *x*_2_(*t*_2_)) (Figure 15, case a). What is the significance of the times *t*_1_ and *t*_2_ with respect to parameter values?
⋄ Do there exist conditions, once the trajectory reaches *V_c_*, under which the optimal trajectory no longer slides? Is it possible that at the time *t*_*_ the point (*x*_1_(*t*_*_), *x*_2_(*t*_*_)) is acontact point (figure 15(b))? Some numerical results suggest that such contact point may exist and give rise to a *YXY* control structure (Figure 12).
⋄ We have shown that an optimal control can switch at most once in each of the regions 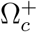 and 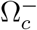. Numerically we did not observe any bang-bang controls of the form *YXY*, although its existence was strongly suggested. On the other hand, Lemma 21 implies that if the optimal control has a switch point when the trajectory lands on region 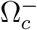 then the optimal trajectory does not slide along *V_c_*. The existence of a bang-bang junction in 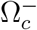 is therefore of interest.
⋄ For all examples plotted in Figures 10 with *d* ≥ 0.1, the entry time occurs approximately at the same value *t*_1_ = 20.03. Is this a coincidence? We would like to understand the dependence of the entry time *t*_1_ and on parameters *α, d, p_r_, M* and/or *ϵ*.
⋄ We would like to extend models to include multiple, possibly non-cross resistant, killing agents. Indeed, clinical practice generally includes multiple agents applied concurrently and sequentially, and we plan on investigating strategies when different types of drugs may be applied. For example, what control strategies arise when a targeted therapy exists which targets the resistant sub-population? What order should the agents be applied, and for how long? Are intermediate doses now optimal? These are all interesting mathematical questions which may be clinically relevant.

**Figure 15:**
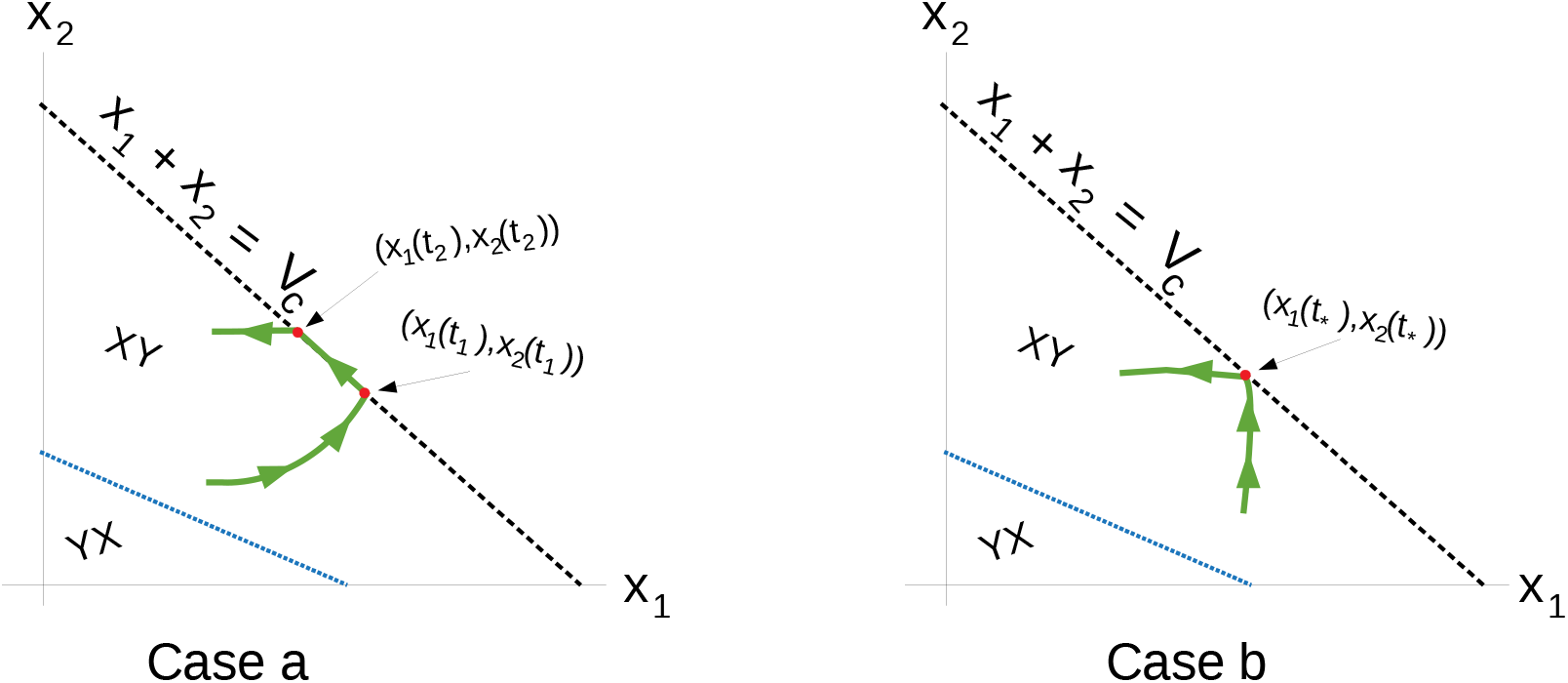
(a) Example of an arc with feedback control where (*x*_1_(*t*_1_), *x*_2_(*t*_1_)) is called the entry point and (*x*_1_(*t*_2_), *x*_2_(*t*_2_)) the exit point (b) Example of an arc that does not slides but reaches the boundary *V* = *V_c_* at the contact point (*x*_1_(*t*_*_), *x*_2_(*t*_*_)).

Drug resistance is one of the main barriers to the success of chemotherapy, and treatment strategies must be designed with this phenomenon in mind. Furthermore, the ability of cancer cells to adapt and express resistance mechanisms in direct response to therapy further complicates the manner in which we administer chemotherapies. In this work, we have attempted to precisely quantify the implications of induced resistance on optimal chemotherapy protocols in an idealized clinical environment where resistance is unavoidable. Induced resistance has been shown to dramatically alter therapy outcome, hence implying that it is vitally important to understand its role in both dynamics and designing chemotherapy regimes.

## 10 Materials and Methods

Numerical solutions were obtained with the use of GPOPS-II software, Invoice number: 0000001256.

## Acknowledgements

This research was supported in part by grant AFOSR FA9550-14-1-0060. We thank Anil Rao for technical suggestions regarding the optimization formulation and the use of GPOPS-II.

## Author Contributions

All authors contributed equally to this work.

## Conflict of Interest

None declared.

## Notes

#### Summary of Updates

Numerical and existence results have been added.

